# Early Seasonal Increases and Persistence in Relative Abundance of Potentially Toxic Cyanobacteria: Concerning Impacts of Extended Ice-Free Periods in Northern Temperate Lakes

**DOI:** 10.1101/2022.12.20.521158

**Authors:** Ellen S. Cameron, Kirsten M. Müller, Mike Stone, Jim Buttle, Jason Leach, Kara Webster, Monica B. Emelko

## Abstract

Cyanobacteria threaten public and ecosystem health globally through the production of secondary metabolites including potent toxins, and disruption of water treatment processes. Warmer water temperatures and high nutrient availability are key characteristics associated with the occurrence of cyanobacteria. There is typically concern of cyanobacteria blooms (e.g., visible biomass accumulations) occurring in the summer season of eutrophic systems. However, in this study, the proliferation of cyanobacteria in lakes across all seasons and in absence of visual biomass indicators of bloom condition was observed in three oligotrophic lakes of the Turkey Lakes Watershed (TLW) in Ontario, located within a sugar maple dominated forest on the Canadian Shield. Almost 40 years of ice phenology data showed that rising temperatures have led to significantly longer ice-free periods and aquatic growing seasons in TLW. Warming is especially evident in the autumn, with the onset of ice-on periods commencing significantly later in the year. Cyanobacterial communities in three interconnected temperate, oligotrophic lakes were characterized over an 18-month period from July 2018 to January 2020 (across 10 synoptic sampling events) using amplicon sequencing of the V4 region of the 16S rRNA gene. During the winter, there was low abundance or occasional absence of cyanobacteria; however, a non-photosynthetic basal lineage of cyanobacteria (Melainabacteria) was present during periods of ice cover. Notably, photosynthetic populations reappeared in the water column immediately following the loss of ice-cover—they were especially abundant in lakes with surficial geology and lake morphometry that favor greater availability of fine sediment and associated nutrients. Thus, this collective analysis demonstrates that the convergence of key abiotic and biotic factors—climate forcing of hydrological and biogeochemical processes, and intrinsic landscape features—enable increases in the relative abundance of potentially toxic cyanobacteria within the temperate forest biome of Canada over increasingly longer periods of time.

## 1. Introduction

Steady increases in harmful algal bloom (HAB) that threaten water and associated public and ecosystem health have been observed in freshwater environments for decades because of increased nutrient availability resulting from anthropogenic (e.g., development, construction of dams/impoundments, river diversion, deforestation, and environmental discharges acting as point and non-point sources) and climate change-exacerbated (e.g., wildfires, extreme precipitation) landscape disturbances (Emelko et al., 2016; Paerl, 2014; Plaas & Paerl, 2021; Winter et al., 2011). It is widely recognized that photosynthetic cyanobacteria threaten water quality and ecosystem health through the formation of dense blooms and the production of secondary metabolites including taste and odour compounds (e.g., geosmin, 2-methyl isoborneol [MIB]) and several cyanotoxins of human and environmental health concern (Harke et al., 2016; Huisman et al., 2018; Vu et al., 2020). These include hepatotoxins, neurotoxins, cytotoxins, and skin and gastrointestinal irritants (Cheung et al., 2013). New toxins and toxin producers also continue to emerge. For example, Melainabacteria are a recently identified non-photosynthetic lineage of cyanobacteria that may produce neurotoxins (Nunes-Costa et al., 2020) that increase the risk of neurodegenerative diseases (Dunlop et al., 2021). They have been observed in municipal wastewater (Lin et al., 2020), lakes (Monchamp et al., 2019a), tap water (Ling et al., 2018), and groundwater (Chik et al., 2020). Critically, the potential drinking water security risks from cyanobacterial proliferation cannot be overstated because most water treatment plants do not have the expensive infrastructure necessary to reliably treat cyanotoxins and taste and odour compounds (Emelko et al., 2011).

Fundamental physiological differences among algal taxa influence their distribution (Irwin et al., 2006, 2012). As growth is typically limited by temperature as well as light and nutrient availability, seasonal and geographic differences in population distributions are common (Butterwick et al., 2005; Dell et al., 2011). In temperate regions, algal populations in lakes are driven by water temperature (Butterwick et al., 2005; Dell et al., 2011) and changes in meteorological conditions that impact light and nutrient availability, and water column stratification (Rusak et al., 2018). While there are exceptions to “typical” seasonal succession in algal populations (Fanesi et al., 2016), in temperate systems: (1) winter communities are typically dominated by cryptophytes, chrysophytes and diatoms, (Beall et al., 2016; Felföldi et al., 2016; Phillips & Fawley, 2002), (2) spring communities include high abundance of diatoms (Jaworska & Zdanowski, 2012; Winder et al., 2009), and (3) summer communities are dominated by green algae (Staehr & Birkeland, 2006; Winder et al., 2009; Winder & Hunter, 2008) and cyanobacteria. Cyanobacterial community composition is further influenced by factors such as nutrient availability (Andersson et al., 2015) with small sized picocyanobacterial (0.2 – 2.0 μm) taxa thriving in low nutrient conditions due to rapid nutrient uptake (Callieri & Stockner, 2002; Collos et al., 2009). Winter limnological processes are relatively poorly understood (Felföldi et al., 2016; Wilhelm et al., 2014) and winter sampling can be logistically complicated because of difficulties in site access, sample collection, and associated safety concerns further perpetuating lack of advancement in winter limnological research. Moreover, lake microbial communities have been traditionally viewed as “dormant” during periods of ice cover (Felföldi et al., 2016; Hampton et al., 2015; Powers & Hampton, 2016). Notably, however, unique winter niches with microbial and phytoplankton communities adapted to low water temperatures and light conditions have been identified (Phillips & Fawley, 2002; Tran et al., 2018) and winter cyanobacterial blooms have been detected during periods of ice cover (Wejnerowski et al., 2018).

Changing climate drives shifts in aquatic microbial community dynamics because it creates environmental conditions that significantly alter interactions between terrestrial and aquatic ecosystems. A North American continental-scale evaluation of thousands of water bodies showed stunning decreases in the number of naturally oligotrophic streams and lakes since the turn of the century because of changing climate and associated landscape disturbances (Stoddard et al., 2016); these changes can be substantial and dominate cumulative watershed effects even after convergence with impacts from anthropogenic disturbances (Watt et al., 2021). A warming climate can alter lake thermal regimes leading to increased water temperature and potentially extended ice-free periods, which may have significant implications for productivity in these aquatic ecosystems (Hampton et al., 2008; O’Beirne et al., 2017). This can alter the timing and magnitude of runoff events from terrestrial to aquatic ecosystems (Creed et al., 2015) and increasing the delivery of nutrients to receiving waters (Kleinman et al., 2011).

The proliferation of cyanobacteria in lakes situated in North American temperate forests has not been systematically characterized, however, especially across all seasons and in absence of extensive visual biomass signaling “bloom” conditions. Such characterization is integral to longer-term evaluation of water quality and treatability implications of climate warming on public health and water security (Emelko et al., 2011) and development of “fit for purpose” adaptation strategies (Blackburn et al., 2021), including nature-based solutions that are frequently cited with “high confidence” as important enablers for reducing adaptation gaps (IPCC, 2022). Here, we characterized cyanobacteria across all seasons in three oligotrophic lakes, located in central Ontario, that have experienced extended ice-free periods. Cyanobacterial community composition was evaluated using amplicon sequencing of the V4 region of the 16S rRNA gene to (1) characterize spatial and seasonal trends, and (2) identify potential treatments to the provision of safe drinking water associated with shifts in the ice-free period in these systems. Seasonal variation in cyanobacterial community composition was evaluated monthly over an 18-month period. Vertical distributions were further evaluated during summer months.

## 2. Methods

### 2.1 TLW Study Site

The TLW Study was established in 1979 to investigate ecosystem effects of acidic atmospheric deposition on terrestrial ecology and surface water quality (Webster et al., 2021). Jeffries et al. (1988) provided a comprehensive description of the physical characteristics of the watershed. In brief, the TLW is situated about 50 km north of Sault Ste. Marie, Ontario on the Canadian Shield in an uneven-aged tolerant hardwood (>90% sugar maple) forest. Geological parent material in the watershed consists of Precambrian silicate greenstone (i.e., metamorphosed basalt) (Jeffries & Semkin, 1983); fine-grained sediment (i.e., glacial till) overlies the bedrock. Four interconnected lakes classified as oligotrophic to mesotrophic are fed by first order streams and groundwater: Batchawana Lake, Wishart Lake, Little Turkey Lake and Big Turkey Lake (Figure S1; Table S1). Except for Wishart, each of the lakes thermally stratifies annually during summer and winter (Figure S2). Wind-induced mixing in this shallow lake generally prevents thermal stratification. During periods of stratification, oxygen is depleted at lower lake depths; zones of anoxia sometimes develop in Batchawana and Little Turkey Lakes. Macrophytes are abundant along the margins of the lakes (Smokorowski et al., 2021) and cyanobacteria were historically identified as dominant members of the phytoplankton communities (Jeffries et al., 1988).

Big Turkey, Little Turkey, and Wishart Lakes were investigated herein. At higher elevations (i.e., Wishart Lake), the till thickness is less than 1 m, with frequent surface exposure of bedrock; there is substantially more till (1 to 2 m) at lower elevation (i.e., Big Turkey Lake, Little Turkey Lake) (Jeffries et al., 1988). As shown in Table S1, the depth ratio (i.e., the ratio of mean to maximum lake depth) is substantially lower in Big Turkey lake than in the other study lakes, as is the renewal time (i.e., how long it takes for inflows to fill the lake).

### 2.2 Ice Phenology Characterization & Hydroclimatic Trends in Turkey Lakes Watershed

Visits by field staff to the vicinity of the TLW study lakes occurred at least weekly and as often as daily during spring melt. Ice status was ascertained from the same viewing location(s) by the same individual during the period of 1980 to 2017. The dates used to characterize ice phenology correspond to permanent, complete ice coverage in the fall and complete disappearance of ice in the spring. The inspection procedure is sufficiently rigorous such that several brief cases of partial and/or temporary ice coverage have been identified. Thus, the dates have an error of at most ± 3 days. Linear regressions were conducted on the ice phenology data (ice on, ice off, total days of ice cover).

Long-term hydroclimatic monitoring data available from the Turkey Lakes Watershed was used to identify trends in changing temperature, precipitation and discharge (Webster et al., 2021). Mean values per month across year were used for linear regression for general long-term trends and local polynomial regression to identify specificities in monthly interannual variability (Figure S2). Data were aggregated into seasonal groupings to examine differences in the ice-free period shoulder seasons, spring (April and May) and autumn (September, October, November) and used for statistical testing.

### 2.3 Sample Collection, DNA Extraction & 16S rRNA Gene Amplicon Sequencing

Water samples were collected and filtered following the approach described in Cameron et al. (2022). They were collected at Secchi depth (during ice-free periods) and 0.25 m below ice (during periods of ice cover), at the deepest locations in Big Turkey, Little Turkey, and Wishart Lakes between July 2018 and January 2020. In July and August of 2018, samples were also collected from the water surface, Secchi depth and one metre below Secchi depth at each lake site to describe the vertical distribution of cyanobacterial communities within the water column, including depths of especially low light intensity that have not been widely investigated. Sampling details are provided in Table S2. As described in Cameron et al. (2022), DNA extraction was performed using the DNeasy PowerSoil Kit (QIAGEN Inc., Venlo, Netherlands), quantified using a NanoDrop spectrophotometer (Table S2; absolute values were only accurate at DNA concentrations of more than 10ng/µL, and submitted to a commercial laboratory (Metagenom Bio Inc.,Waterloo, ON) for amplicon sequencing of the V4 region of the 16S rRNA gene (515FB:[GTGYCAGCMGCCGCGGTAA]; 806RB – [GGACTACNVGGGTWTCTAAT]; Walters et al., 2015) sequencing using the Illumina MiSeq platform (Illumina Inc., San Diego, United States).

### 2.4 Sequence Processing & Community Analyses

Amplicon sequencing data were processed and normalized using the approaches described in Cameron et al. (2021). In brief, bioinformatic processing was conducted using QIIME2 (v. 2019.10; Bolyen et al., 2019). The amplicon sequence variant (ASV) table was constructed using DADA2 (Callahan et al., 2016) and ASVs were taxonomically classified using a Naïve-Bayes taxonomic classifier trained using the SILVA138 database (Quast et al., 2013; Yilmaz et al., 2014). Sequences from the 16S rRNA gene classified as mitochondria or chloroplasts were removed to centralize focus to bacterial community members. Taxonomic classifications for sequence variants classified as Cyanobacteria at the phylum level were manually curated to reflect taxonomic levels according to AlgaeBase (Guiry & Guiry, 2022) and confirmed using a phylogenetic tree constructed using Cydrasil (Roush et al., 2021) and visualized with Interactive tree of life (iTOL; Letunic & Bork, 2016; Figure S5; Table S6). Downstream analyses were conducted in R (v. 4.0.) utilizing *qiime2R* (v. 0.99.23; Bisanz, 2018) for import of QIIME2 files into the R environment, and *phyloseq* (v. 1.32.0; McMurdie & Holmes, 2012) for storage and handling of processed sequencing data. For cyanobacterial community analysis, ASVs classified as Cyanobacteria at the phylum level were filtered to create libraries consisting of only cyanobacteria classified sequences. Cyanobacterial libraries were repeatedly rarefied to a normalized size of 824 reads using *mirlyn* (Cameron & Tremblay, 2020) to address challenges of amplicon sequencing partially representing source diversity (Schmidt et al., 2022).

The composition of cyanobacterial communities was assessed at the taxonomic order level, and relative abundances were visualized using a heatmap. Relative abundances were randomized across phyla within samples to identify significantly dominant groups within the bacterial community in relation to sampling conditions with a Bonferroni correction. Cyanobacterial community composition was visualized at the order level using *ComplexHeatMap* (v. 2.12.1; Gu et al., 2016). Cyanobacterial orders present at <5% relative abundance cumulatively across samples were excluded from composition visualization. To further examine the composition of cyanobacterial communities, relative abundances of specific ASVs in the cyanobacterial community classified to the order Synechococcales were plotted to demonstrate the potential dominance of individual ASVs within the community.

The Shannon Index (Shannon, 1948) was used to identify within sample trends in alpha diversity as a function of sampling depth or sampling month. To characterize similarity in community composition, rarefied libraries were used to calculate Bray-Curtis distances (Bray & Curtis, 1957) and subsequently average values were displayed using a heatmap generated with *ComplexHeatMap*.

### 2.5 Data Availability

Sequencing data analyzed in this study are available in the European Nucleotide Archive (ENA) under study accession PRJEB56927.

## 3. Results

### 3.1 Evidence of Extension of the Ice-Free Period & Seasonal Differences in Shoulder Seasons

Evidence of changes in ice phenology (date of permanent, complete ice coverage; date of complete disappearance of ice; and days of complete ice cover) in the TLW during the period from 1980 to 2017 are presented in Figure 1. Significant changes in the date of ice disappearance were not observed in any of the lakes (*p* > 0.25). The date of complete ice disappearance ranged from March to May in each of the lakes; specifically, Julian day 88 (March 29) to 140 (May 20) in Big Turkey Lake, day 88 (March 29) to 140 (May 20) in both Little Turkey and Wishart Lakes. In contrast, changes in the date of ice cover onset were significant in each of the lakes (p = 0.012, 0.012, and 0.006, respectively), indicating a trend of fall ice formation occurring later in the year (Figure 1), likely because of increasing air temperature in the region. The date of initiation of complete ice cover ranged from Julian day 319 (November 15) to 365 (December 31) in Big Turkey Lake, day 316 (November 12) to 365 (December 31) in Little Turkey Lake, and day 309 (November 5) to 356 (December 22) in Wishart Lake. These changes resulted in a significant decline (-0.61, -0.53, and -0.56 days/year, respectively) in the annual period of ice cover in each of the lakes (*p* = 0.021, 0.053, 0.042, respectively).

**Figure 1:**
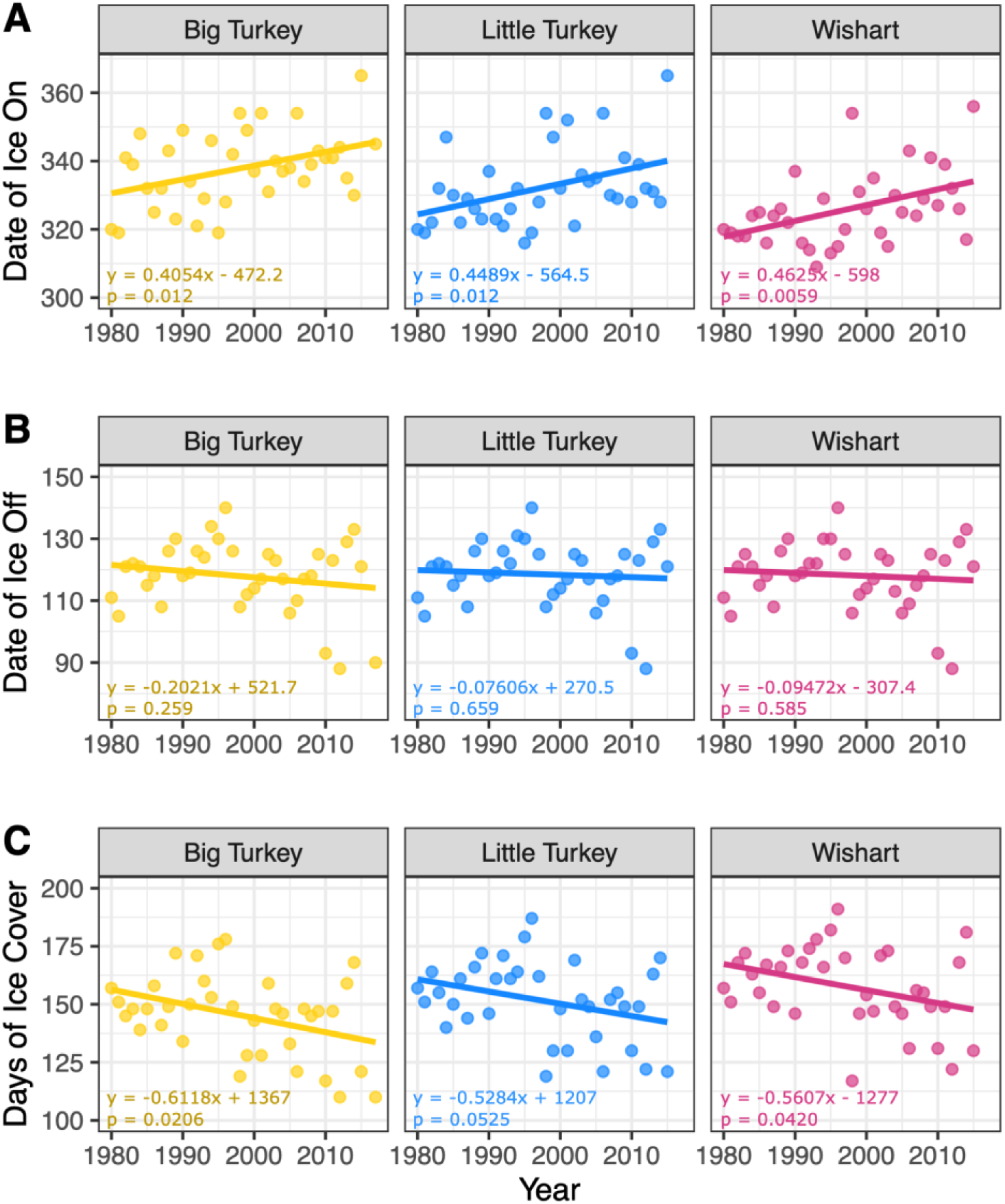
Ice phenology of the lakes of Turkey Lakes Watershed from 1980 – 2015 demonstrating the (A) date of ice formation (Julian calendar day number), (B) date of ice loss (Juan calendar day number) and (C) total days of ice cover annually. Slopes for date of ice on and annual days of ice cover were significant (p < 0.05) indicating significant lengthening of ice-free periods that extend to a later date annually.

Examination of the aggregated ice-free shoulder seasons (*i.e.*, spring [April-May] and autumn [September-October-November]) in the TLW revealed key climatic differences expected to impact the physicochemical conditions of the lakes.

Corresponding to ice cover loss and subsequent snowmelt, increased discharge in the watershed was observed in April and May (Figure 2). Specifically, while the autumn had significantly higher (p = 3.761e-14) precipitation (μ = 138.68 *mm*) than the spring (μ = 79.81 *mm*), discharge in the spring (*μ* = 128.27 *mm*) was significantly (p = 1.772e-06) higher (μ = 74.08 *mm)* in the autumn. Mean air temperatures were observed to be significantly higher (p = 5.063e-09) in the spring (μ = 7.16°C) than the autumn (μ = 3.10°C). While air temperatures do not directly equate to water temperatures, values as these can be used to inform the expected trends observed in lake systems. Available data on water column stratification (Figure S2) reveals surface water temperature of lakes warms rapidly following loss of ice cover in the spring (e.g., May 2019). Surface water temperatures greater than 20°C were observed in lakes as quickly as 4 weeks following ice-cover loss due to the rapid warming as a result of increased air temperatures and increased solar irradiance.

**Figure 2:**
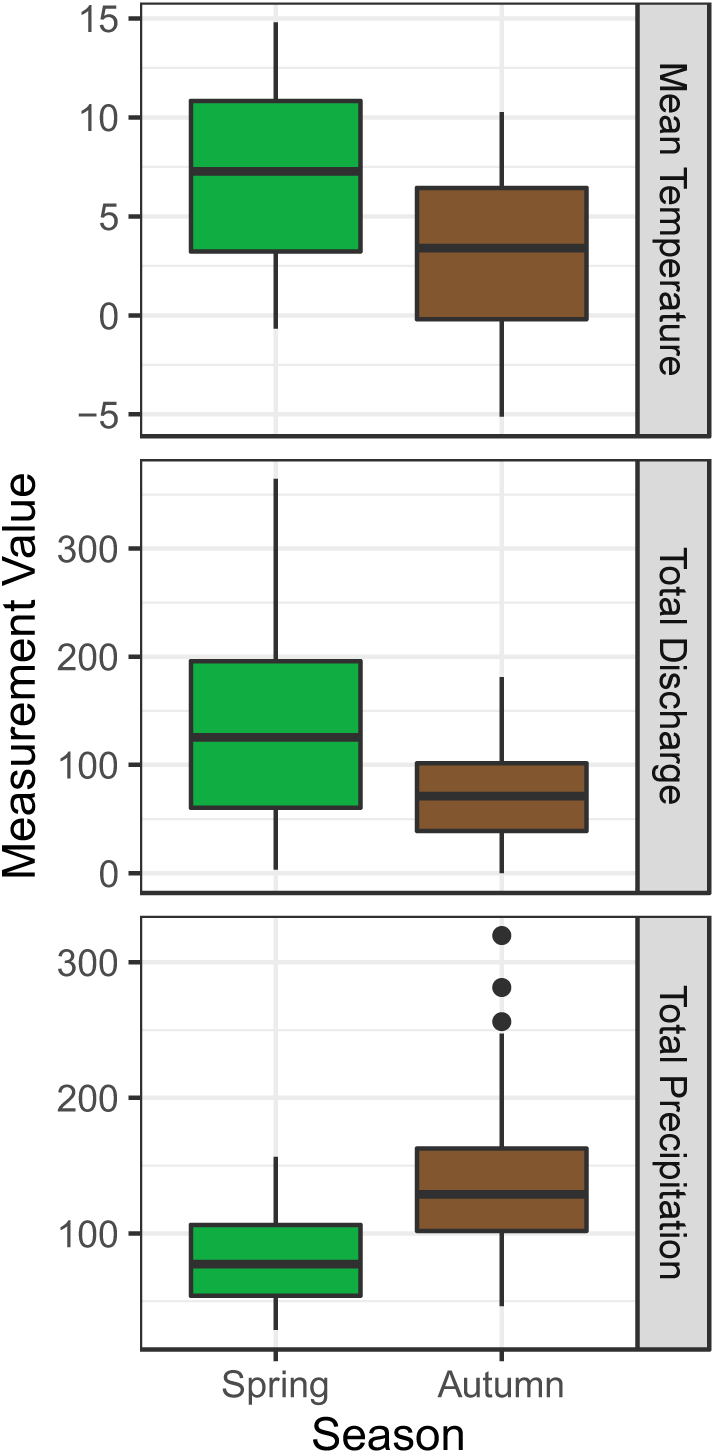
Seasonal aggregations of hydroclimatic data reflecting trends in the mean monthly temperature (°C), total discharge (mm) and total precipitation (mm). Significant differences exist between both seasons across all three measures potentially driving cyanobacterial population dynamics.

### 3.2 Cyanobacteria Populations Persist Year-Round and Fluctuate Seasonally During Ice-Free Periods

Cyanobacterial 16S rRNA gene sequences were observed in samples from at least some of the TLW study lakes on every sampling occasion including periods of ice cover (Table 1), though their relative abundances in the bacterial community fluctuated seasonally (0 to 56.3%; Figure 3; Table 1). Observed cyanobacterial amplicon sequence variants were classified to potentially toxic, bloom forming taxa belonging to the orders Chroococcales, Nostocales and Synechococcales (Table 3) including *Microcystis*, *Pseudanabaena*, *Anabaena*, and *Aphanizomenon*. In Big Turkey Lake, cyanobacterial sequences were significantly represented in the bacterial community during all ice-free sampling months (*p* < 0.05; Table S4; Figure S6) except August 2018 (*p* = 0.1). In Little Turkey and Wishart Lakes, they were significantly represented in the bacterial communities during the summer months (June, July, August) (p < 0.009), except August 2018 (*p* = 1). However, unlike bacterial communities in Big Turkey Lake, cyanobacterial sequences were not significantly represented in October and May in either Little Turkey or Wishart Lake (*p* = 1). Although they were observed during periods of ice cover (February 2019, March 2019, and January 2020), the relative abundance of cyanobacterial sequences was often low; thus, cyanobacteria were not significantly represented in any of the study lakes although they were often present (Table S4; Figure S6; Turkey *p* > 0.96; Little Turkey *p* = 1; Wishart *p* = 1). During these months, cyanobacteria represented less than 1% of the total bacterial community and the detected sequences were largely composed of variants classified to the under-studied non-photosynthetic basal lineages of cyanobacteria: Melainabacteria and Sericytochromatia (Figure S4; Table 3).

**Figure 3:**
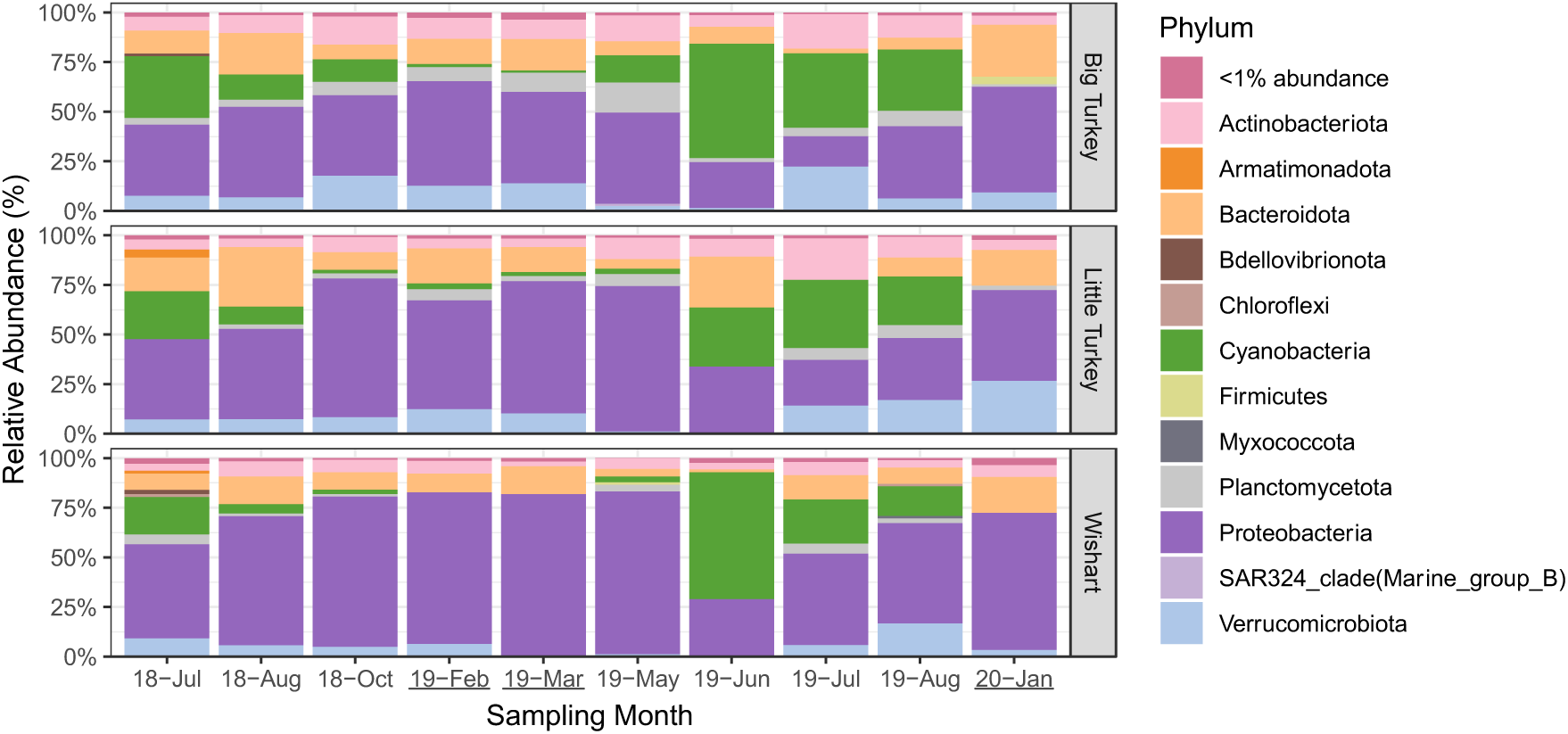
Stacked bar charts depicting the relative abundances of major bacterial phyla identified from amplicon sequencing of the V4 region of the 16S rRNA gene across a multi-seasonal timeframe in a non-stratified lake (Wishart), a mid-sized stratified lake (Little Turkey) and deep stratified lake (Big Turkey). Underlined sampling months indicate periods of ice cover.

**Table 1.**
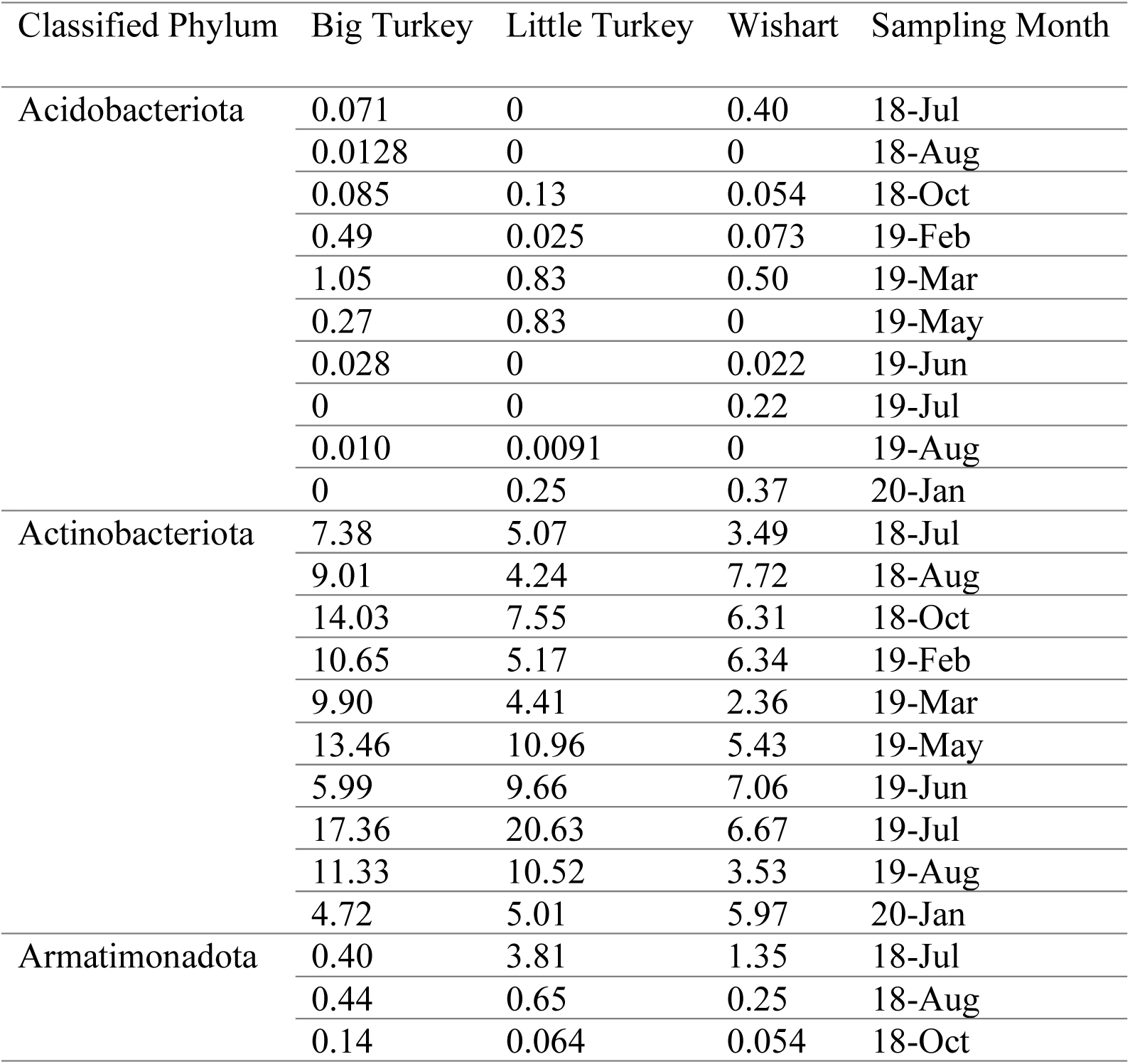
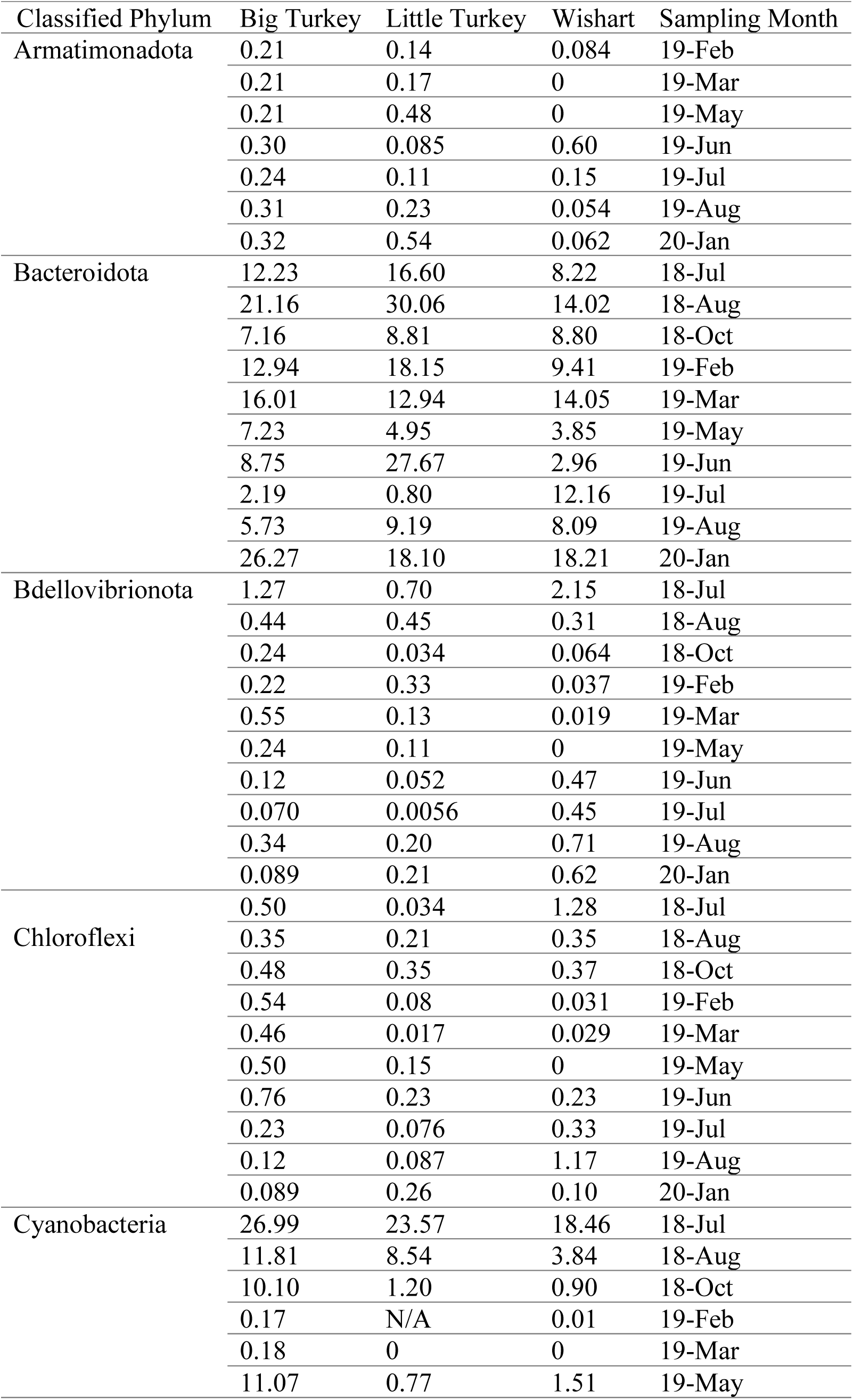
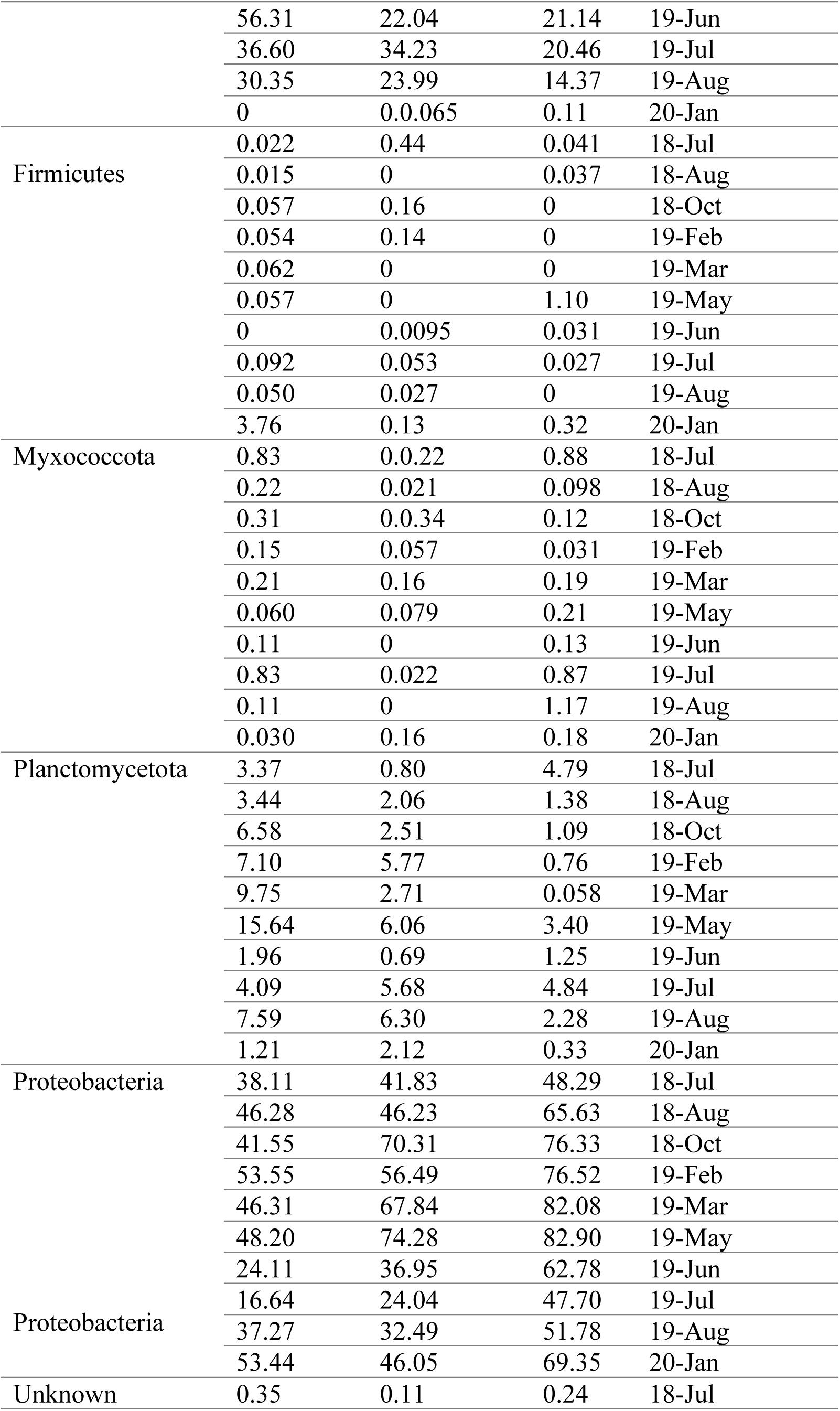
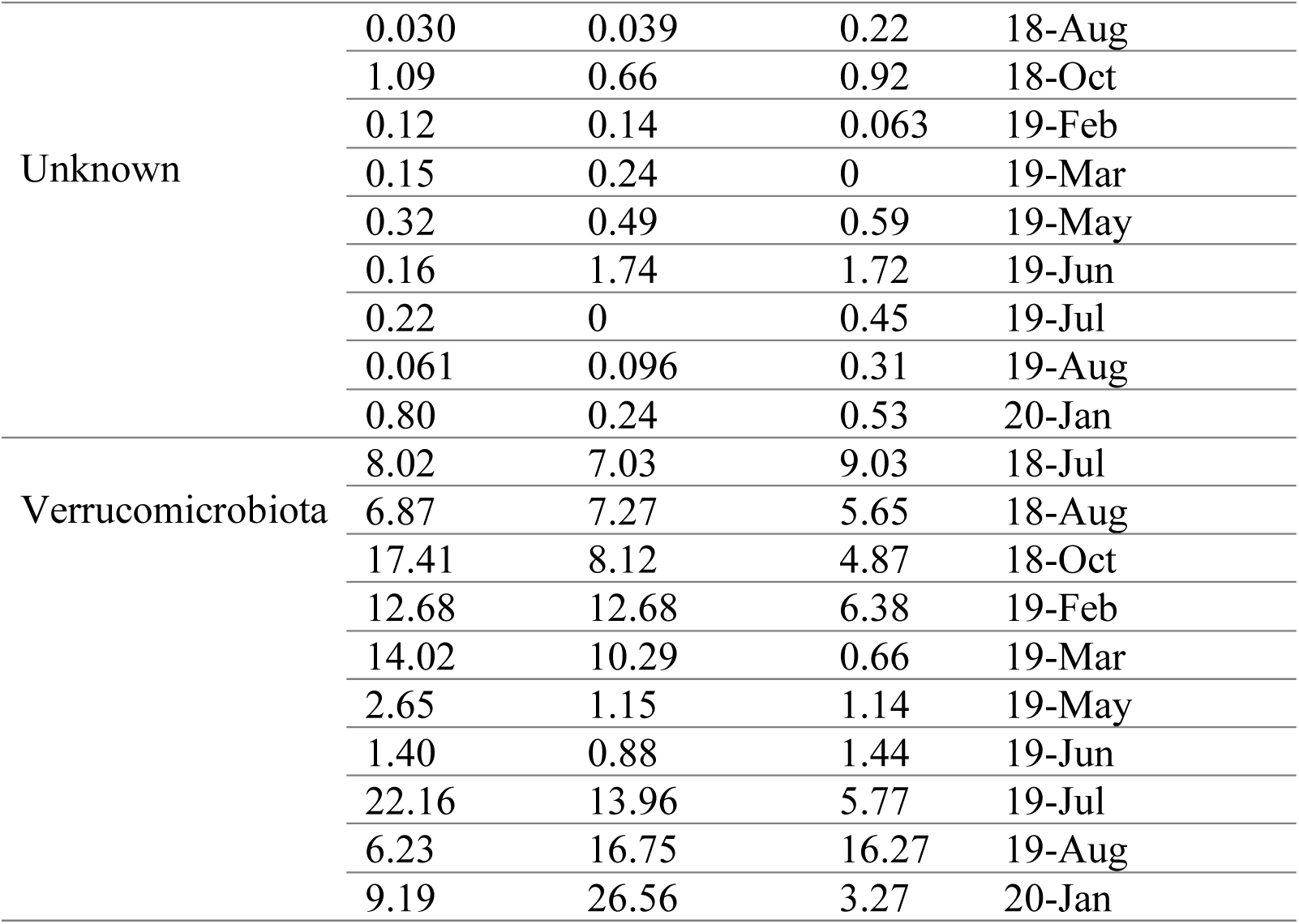
Relative abundances of major bacterial phyla rounded to two decimal points across a multi-seasonal profile. Phyla present at less than 1% abundance across all samples were excluded from this table.

| Classified Phylum | Big Turkey | Little Turkey | Wishart | Sampling Month |
| --- | --- | --- | --- | --- |
| Acidobacteriota | 0.071 | 0 | 0.40 | 18-Jul |
|  | 0.0128 | 0 | 0 | 18-Aug |
|  | 0.085 | 0.13 | 0.054 | 18-Oct |
|  | 0.49 | 0.025 | 0.073 | 19-Feb |
|  | 1.05 | 0.83 | 0.50 | 19-Mar |
|  | 0.27 | 0.83 | 0 | 19-May |
|  | 0.028 | 0 | 0.022 | 19-Jun |
|  | 0 | 0 | 0.22 | 19-Jul |
|  | 0.010 | 0.0091 | 0 | 19-Aug |
|  | 0 | 0.25 | 0.37 | 20-Jan |
| Actinobacteriota | 7.38 | 5.07 | 3.49 | 18-Jul |
|  | 9.01 | 4.24 | 7.72 | 18-Aug |
|  | 14.03 | 7.55 | 6.31 | 18-Oct |
|  | 10.65 | 5.17 | 6.34 | 19-Feb |
|  | 9.90 | 4.41 | 2.36 | 19-Mar |
|  | 13.46 | 10.96 | 5.43 | 19-May |
|  | 5.99 | 9.66 | 7.06 | 19-Jun |
|  | 17.36 | 20.63 | 6.67 | 19-Jul |
|  | 11.33 | 10.52 | 3.53 | 19-Aug |
|  | 4.72 | 5.01 | 5.97 | 20-Jan |
| Armatimonadota | 0.40 | 3.81 | 1.35 | 18-Jul |
|  | 0.44 | 0.65 | 0.25 | 18-Aug |
|  | 0.14 | 0.064 | 0.054 | 18-Oct |

| **Table 1 Continued** |  |  |  |  |
| --- | --- | --- | --- | --- |
| Classified Phylum | Big Turkey | Little Turkey | Wishart | Sampling Month |
| Armatimonadota | 0.21 | 0.14 | 0.084 | 19-Feb |
|  | 0.21 | 0.17 | 0 | 19-Mar |
|  | 0.21 | 0.48 | 0 | 19-May |
|  | 0.30 | 0.085 | 0.60 | 19-Jun |
|  | 0.24 | 0.11 | 0.15 | 19-Jul |
|  | 0.31 | 0.23 | 0.054 | 19-Aug |
|  | 0.32 | 0.54 | 0.062 | 20-Jan |
| Bacteroidota | 12.23 | 16.60 | 8.22 | 18-Jul |
|  | 21.16 | 30.06 | 14.02 | 18-Aug |
|  | 7.16 | 8.81 | 8.80 | 18-Oct |
|  | 12.94 | 18.15 | 9.41 | 19-Feb |
|  | 16.01 | 12.94 | 14.05 | 19-Mar |
|  | 7.23 | 4.95 | 3.85 | 19-May |
|  | 8.75 | 27.67 | 2.96 | 19-Jun |
|  | 2.19 | 0.80 | 12.16 | 19-Jul |
|  | 5.73 | 9.19 | 8.09 | 19-Aug |
|  | 26.27 | 18.10 | 18.21 | 20-Jan |
| Bdellovibrionota | 1.27 | 0.70 | 2.15 | 18-Jul |
|  | 0.44 | 0.45 | 0.31 | 18-Aug |
|  | 0.24 | 0.034 | 0.064 | 18-Oct |
|  | 0.22 | 0.33 | 0.037 | 19-Feb |
|  | 0.55 | 0.13 | 0.019 | 19-Mar |
|  | 0.24 | 0.11 | 0 | 19-May |
|  | 0.12 | 0.052 | 0.47 | 19-Jun |
|  | 0.070 | 0.0056 | 0.45 | 19-Jul |
|  | 0.34 | 0.20 | 0.71 | 19-Aug |
|  | 0.089 | 0.21 | 0.62 | 20-Jan |
| Chloroflexi | 0.50 | 0.034 | 1.28 | 18-Jul |
|  | 0.35 | 0.21 | 0.35 | 18-Aug |
|  | 0.48 | 0.35 | 0.37 | 18-Oct |
|  | 0.54 | 0.08 | 0.031 | 19-Feb |
|  | 0.46 | 0.017 | 0.029 | 19-Mar |
|  | 0.50 | 0.15 | 0 | 19-May |
|  | 0.76 | 0.23 | 0.23 | 19-Jun |
|  | 0.23 | 0.076 | 0.33 | 19-Jul |
|  | 0.12 | 0.087 | 1.17 | 19-Aug |
|  | 0.089 | 0.26 | 0.10 | 20-Jan |
| Cyanobacteria | 26.99 | 23.57 | 18.46 | 18-Jul |
|  | 11.81 | 8.54 | 3.84 | 18-Aug |
|  | 10.10 | 1.20 | 0.90 | 18-Oct |
|  | 0.17 | N/A | 0.01 | 19-Feb |
|  | 0.18 | 0 | 0 | 19-Mar |
|  | 11.07 | 0.77 | 1.51 | 19-May |
|  | 56.31 | 22.04 | 21.14 | 19-Jun |
|  | 36.60 | 34.23 | 20.46 | 19-Jul |
|  | 30.35 | 23.99 | 14.37 | 19-Aug |
|  | 0 | 0.0065 | 0.11 | 20-Jan |
| Firmicutes | 0.022 | 0.44 | 0.041 | 18-Jul |
|  | 0.015 | 0 | 0.037 | 18-Aug |
|  | 0.057 | 0.16 | 0 | 18-Oct |
|  | 0.054 | 0.14 | 0 | 19-Feb |
|  | 0.062 | 0 | 0 | 19-Mar |
|  | 0.057 | 0 | 1.10 | 19-May |
|  | 0 | 0.0095 | 0.031 | 19-Jun |
|  | 0.092 | 0.053 | 0.027 | 19-Jul |
|  | 0.050 | 0.027 | 0 | 19-Aug |
|  | 3.76 | 0.13 | 0.32 | 20-Jan |
| Myxococcota | 0.83 | 0.022 | 0.88 | 18-Jul |
|  | 0.22 | 0.021 | 0.098 | 18-Aug |
|  | 0.31 | 0.034 | 0.12 | 18-Oct |
|  | 0.15 | 0.057 | 0.031 | 19-Feb |
|  | 0.21 | 0.16 | 0.19 | 19-Mar |
|  | 0.060 | 0.079 | 0.21 | 19-May |
|  | 0.11 | 0 | 0.13 | 19-Jun |
|  | 0.83 | 0.022 | 0.87 | 19-Jul |
|  | 0.11 | 0 | 1.17 | 19-Aug |
|  | 0.030 | 0.16 | 0.18 | 20-Jan |
| Planctomycetota | 3.37 | 0.80 | 4.79 | 18-Jul |
|  | 3.44 | 2.06 | 1.38 | 18-Aug |
|  | 6.58 | 2.51 | 1.09 | 18-Oct |
|  | 7.10 | 5.77 | 0.76 | 19-Feb |
|  | 9.75 | 2.71 | 0.058 | 19-Mar |
|  | 15.64 | 6.06 | 3.40 | 19-May |
|  | 1.96 | 0.69 | 1.25 | 19-Jun |
|  | 4.09 | 5.68 | 4.84 | 19-Jul |
|  | 7.59 | 6.30 | 2.28 | 19-Aug |
|  | 1.21 | 2.12 | 0.33 | 20-Jan |
| Proteobacteria | 38.11 | 41.83 | 48.29 | 18-Jul |
|  | 46.28 | 46.23 | 65.63 | 18-Aug |
|  | 41.55 | 70.31 | 76.33 | 18-Oct |
|  | 53.55 | 56.49 | 76.52 | 19-Feb |
|  | 46.31 | 67.84 | 82.08 | 19-Mar |
|  | 48.20 | 74.28 | 82.90 | 19-May |
|  | 24.11 | 36.95 | 62.78 | 19-Jun |
| Proteobacteria | 16.64 | 24.04 | 47.70 | 19-Jul |
|  | 37.27 | 32.49 | 51.78 | 19-Aug |
|  | 53.44 | 46.05 | 69.35 | 20-Jan |
| Unknown | 0.35 | 0.11 | 0.24 | 18-Jul |
| Unknown | 0.030 | 0.039 | 0.22 | 18-Aug |
|  | 1.09 | 0.66 | 0.92 | 18-Oct |
|  | 0.12 | 0.14 | 0.063 | 19-Feb |
|  | 0.15 | 0.24 | 0 | 19-Mar |
|  | 0.32 | 0.49 | 0.59 | 19-May |
|  | 0.16 | 1.74 | 1.72 | 19-Jun |
|  | 0.22 | 0 | 0.45 | 19-Jul |
|  | 0.061 | 0.096 | 0.31 | 19-Aug |
|  | 0.80 | 0.24 | 0.53 | 20-Jan |
| Verrucomicrobiota | 8.02 | 7.03 | 9.03 | 18-Jul |
|  | 6.87 | 7.27 | 5.65 | 18-Aug |
|  | 17.41 | 8.12 | 4.87 | 18-Oct |
|  | 12.68 | 12.68 | 6.38 | 19-Feb |
|  | 14.02 | 10.29 | 0.66 | 19-Mar |
|  | 2.65 | 1.15 | 1.14 | 19-May |
|  | 1.40 | 0.88 | 1.44 | 19-Jun |
|  | 22.16 | 13.96 | 5.77 | 19-Jul |
|  | 6.23 | 16.75 | 16.27 | 19-Aug |
|  | 9.19 | 26.56 | 3.27 | 20-Jan |

### 3.3 Picocyanobacteria Dominate Cyanobacterial Communities During Ice-Free Periods

Sequences classified to the order Synechococcales (84.4 to 99.2%; n = 74 ASVs) consistently dominated cyanobacterial communities across sampling depths (Figure 4; Table 4) and seasonally (69.0 to 99.8%; Figure 5 Table 1; n = 49) during ice-free periods. Other orders composed the remainder of the cyanobacterial community (Nostocales <1 to 2.3%; Table 3,4; Chroococcales 2.4 to 30.8%). The higher abundances of Chroococcales, the order containing toxic bloom-forming genera such as *Microcystis*, were notably observed at highest abundances in the late summer and shallower sampling depths.

**Figure 4:**
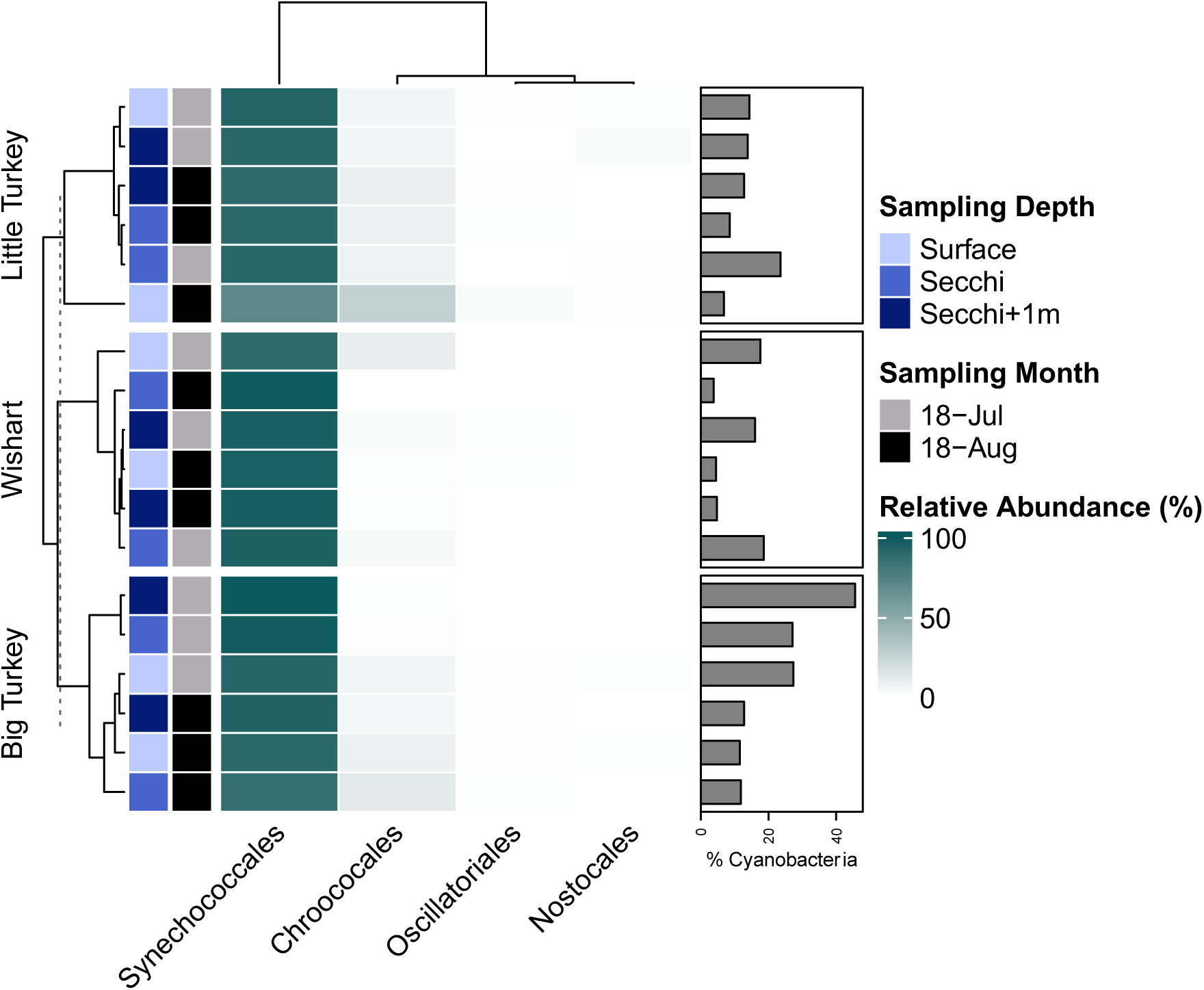
Taxonomic composition of cyanobacterial communities at the order level across a water column profile to identify spatial distribution of different cyanobacterial populations. Relative abundances of cyanobacterial orders are visualized in the heatmap with decreasing darkness representative of higher abundances. Overall proportion of cyanobacterial sequences in the bacterial community are visualized in the right hand side bar graph to provide community composition contextualization. Cyanobacterial orders present in low abundance (<5% cumulatively across all samples) were excluded from visualization.

**Figure 5:**
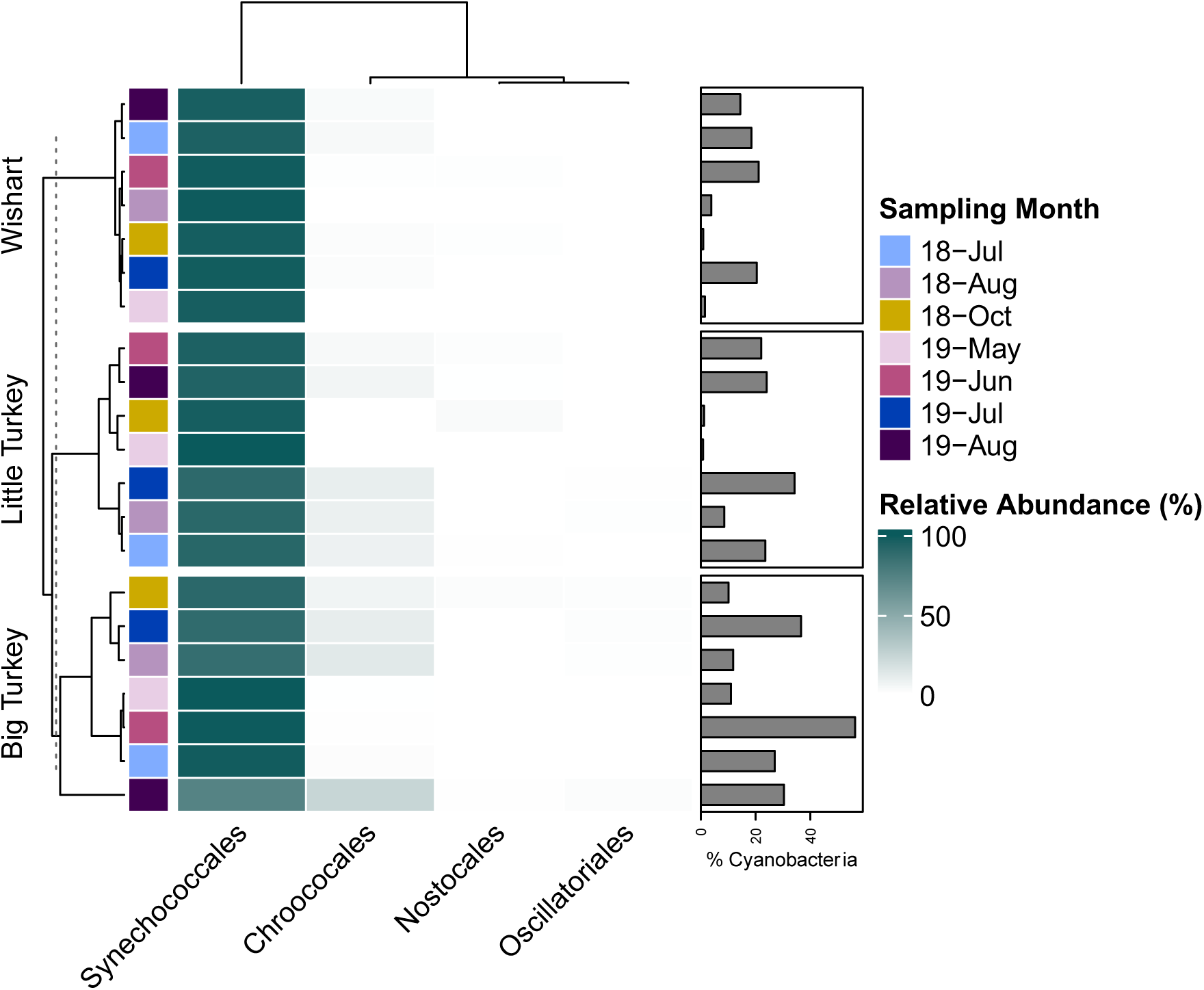
Taxonomic composition of cyanobacterial communities at the order level across a long-term seasonal profile to identify trends in seasonal succession. Relative abundances of cyanobacterial orders are visualized in the heatmap with decreasing darkness representative of higher abundances. Overall proportion of cyanobacterial sequences in the bacterial community are visualized in the right hand side bar graph to provide community composition contextualization. Cyanobacterial orders present in low abundance (<5% cumulatively across all samples) were excluded from visualization.

**Table 2.**
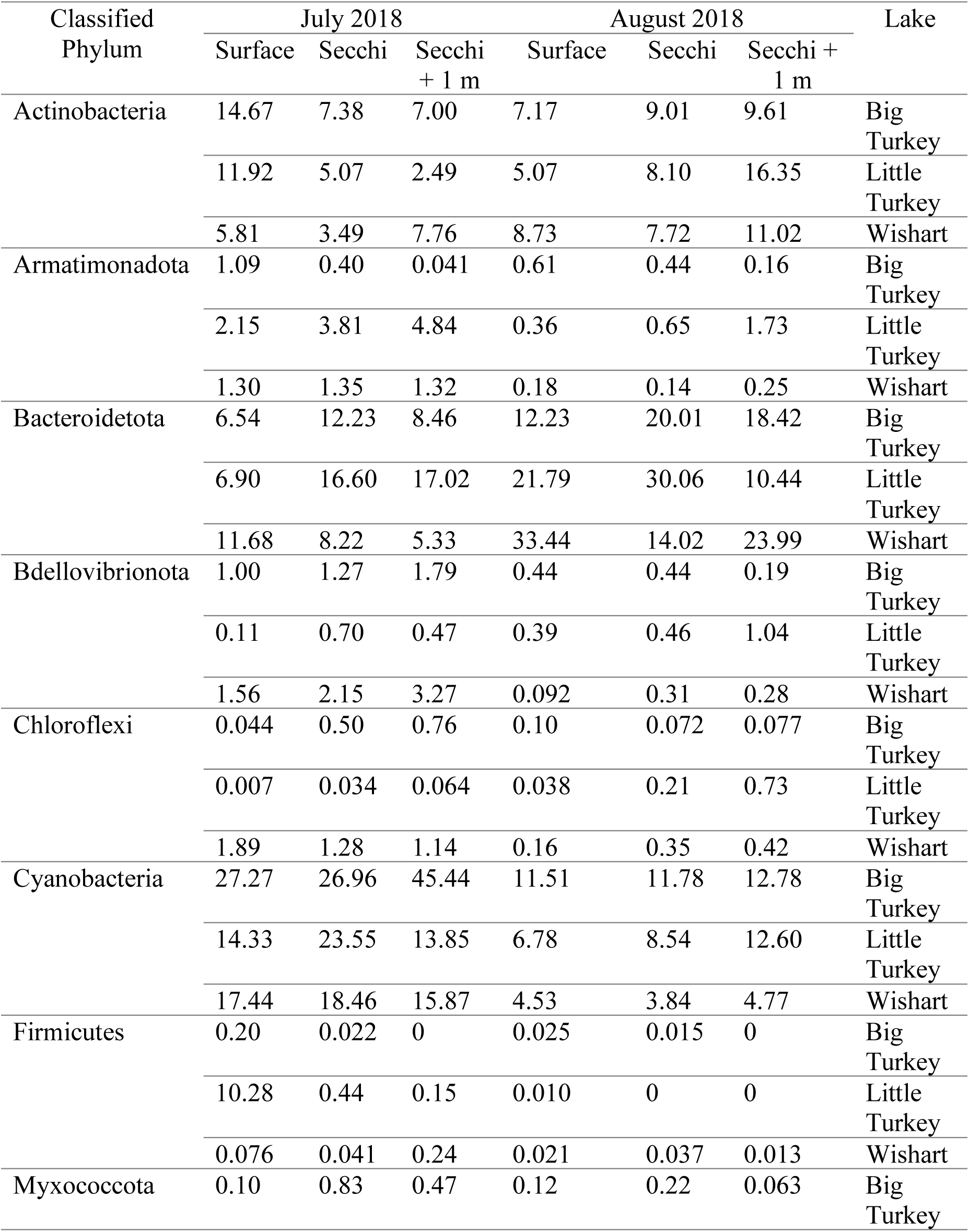
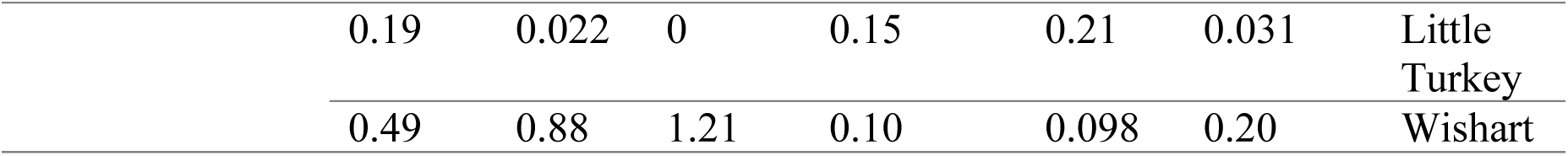
Relative abundances of major bacterial phyla rounded to two decimal points across a depth profile in the summer of 2018. Phyla present at less than 1% abundance across all samples were excluded from this table.

| Classified<br>Phylum | July 2018 |  |  | August 2018 |  |  | Lake |
| --- | --- | --- | --- | --- | --- | --- | --- |
|  | Surface | Secchi | Secchi<br>+ 1 m | Surface | Secchi | Secchi +<br>1 m |  |
| Actinobacteria | 14.67 | 7.38 | 7.00 | 7.17 | 9.01 | 9.61 | Big<br>Turkey |
|  | 11.92 | 5.07 | 2.49 | 5.07 | 8.10 | 16.35 | Little<br>Turkey |
|  | 5.81 | 3.49 | 7.76 | 8.73 | 7.72 | 11.02 | Wishart |
| Armatimonadota | 1.09 | 0.40 | 0.041 | 0.61 | 0.44 | 0.16 | Big<br>Turkey |
|  | 2.15 | 3.81 | 4.84 | 0.36 | 0.65 | 1.73 | Little<br>Turkey |
|  | 1.30 | 1.35 | 1.32 | 0.18 | 0.14 | 0.25 | Wishart |
| Bacteroidetota | 6.54 | 12.23 | 8.46 | 12.23 | 20.01 | 18.42 | Big<br>Turkey |
|  | 6.90 | 16.60 | 17.02 | 21.79 | 30.06 | 10.44 | Little<br>Turkey |
|  | 11.68 | 8.22 | 5.33 | 33.44 | 14.02 | 23.99 | Wishart |
| Bdellovibrionota | 1.00 | 1.27 | 1.79 | 0.44 | 0.44 | 0.19 | Big<br>Turkey |
|  | 0.11 | 0.70 | 0.47 | 0.39 | 0.46 | 1.04 | Little<br>Turkey |
|  | 1.56 | 2.15 | 3.27 | 0.092 | 0.31 | 0.28 | Wishart |
| Chloroflexi | 0.044 | 0.50 | 0.76 | 0.10 | 0.072 | 0.077 | Big<br>Turkey |
|  | 0.007 | 0.034 | 0.064 | 0.038 | 0.21 | 0.73 | Little<br>Turkey |
|  | 1.89 | 1.28 | 1.14 | 0.16 | 0.35 | 0.42 | Wishart |
| Cyanobacteria | 27.27 | 26.96 | 45.44 | 11.51 | 11.78 | 12.78 | Big<br>Turkey |
|  | 14.33 | 23.55 | 13.85 | 6.78 | 8.54 | 12.60 | Little<br>Turkey |
|  | 17.44 | 18.46 | 15.87 | 4.53 | 3.84 | 4.77 | Wishart |
| Firmicutes | 0.20 | 0.022 | 0 | 0.025 | 0.015 | 0 | Big<br>Turkey |
|  | 10.28 | 0.44 | 0.15 | 0.010 | 0 | 0 | Little<br>Turkey |
|  | 0.076 | 0.041 | 0.24 | 0.021 | 0.037 | 0.013 | Wishart |
| Myxococcota | 0.10 | 0.83 | 0.47 | 0.12 | 0.22 | 0.063 | Big<br>Turkey |
|  | 0.19 | 0.022 | 0 | 0.15 | 0.21 | 0.031 | Little<br>Turkey |
|  | 0.49 | 0.88 | 1.21 | 0.10 | 0.098 | 0.20 | Wishart |

**Table 3:**
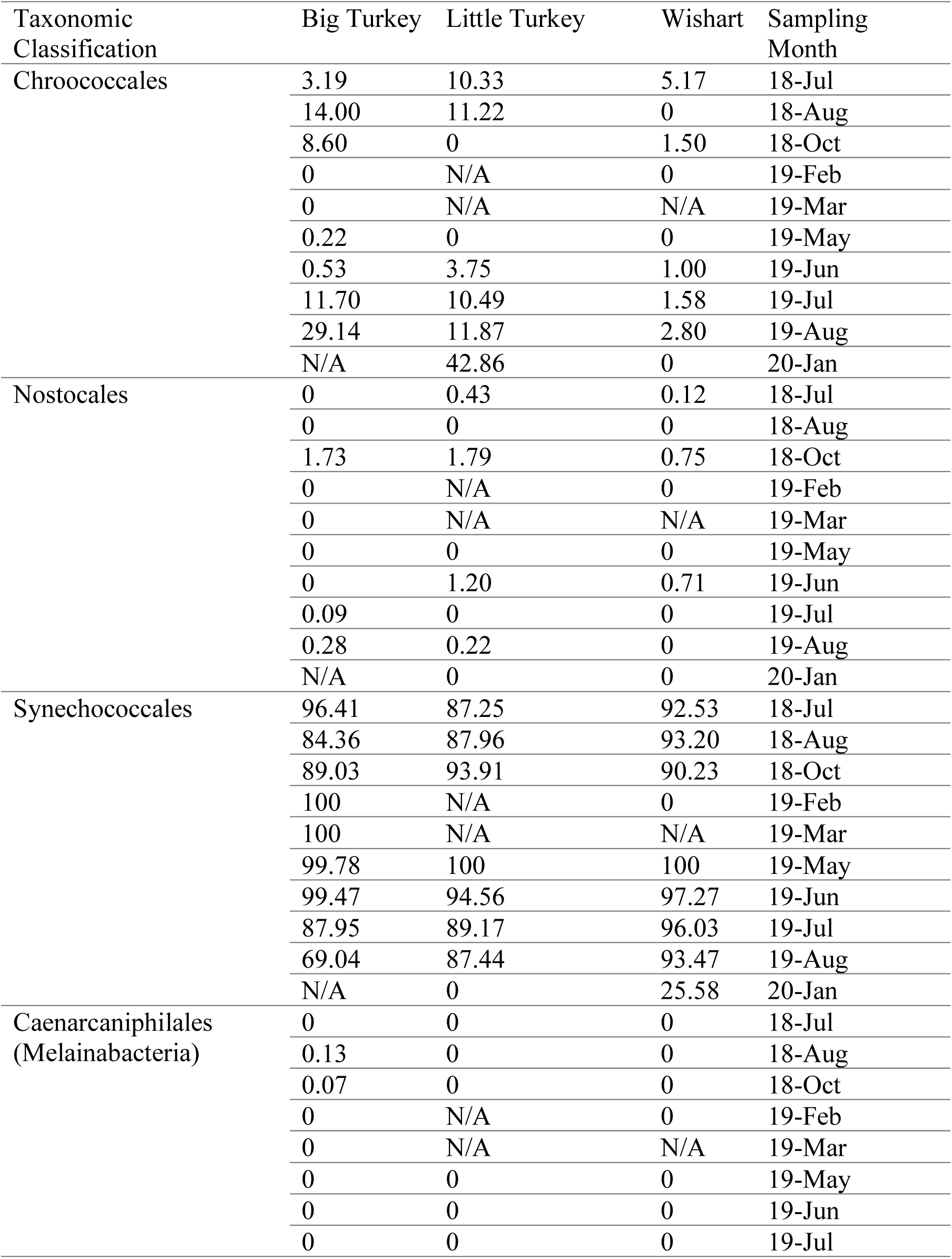
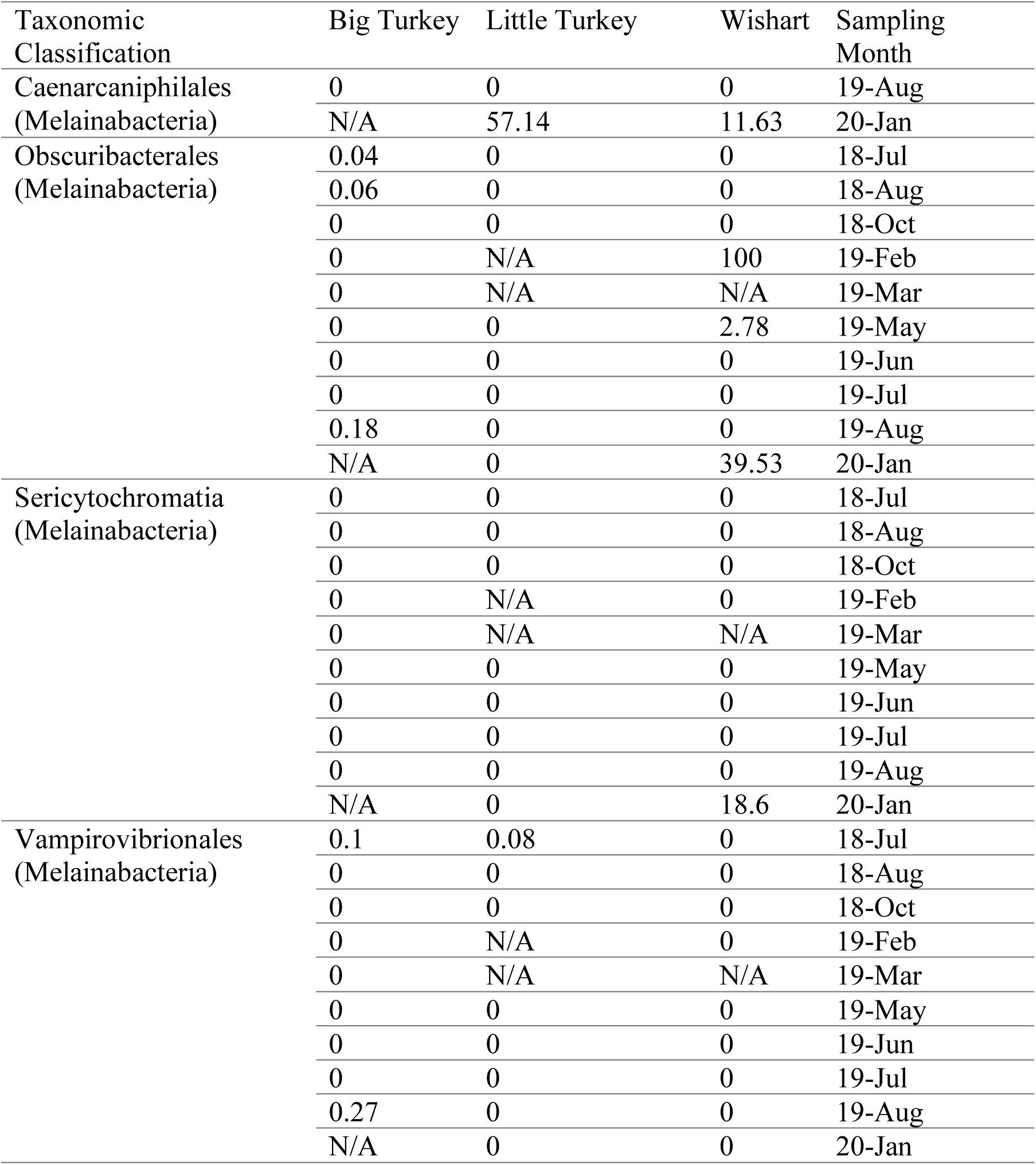
Relative abundances of sequences classified to cyanobacterial taxonomic orders that compose the cyanobacterial community across a seasonal period.

**Table 4:** Relative abundances of sequences classified to cyanobacterial taxonomic orders that compose the cyanobacterial community across a depth profile.

| Taxonomic<br>Clasification | July 2018 |  |  | August 2018 |  |  | Sampling<br>Depth |
| --- | --- | --- | --- | --- | --- | --- | --- |
|  | Big<br>Turkey | Little<br>Turkey | Wishart | Big<br>Turkey | Little<br>Turkey | Wishart |  |
| Chroococcales | 6.96 | 9.13 | 12.04 | 9.28 | 30.76 | 3.30 | Surface |
|  | 3.20 | 10.34 | 5.18 | 14.00 | 11.22 | 0 | Secchi |
|  | 0.75 | 6.26 | 5.69 | 6.98 | 12.22 | 2.43 | Secchi +<br>1 m |
| Nostocales | 1.38 | 0.68 | 0 | 0 | 0 | 0 | Surface |
|  | 0 | 0.43 | 0.12 | 0.75 | 0 | 0 | Secchi |
|  | 0 | 2.31 | 0 | 0 | 0 | 0 | Secchi +<br>1 m |
| Synechococcales | 90.16 | 86.65 | 86.48 | 88.97 | 65.18 | 89.39 | Surface |
|  | 96.41 | 87.26 | 92.53 | 84.37 | 87.96 | 93.20 | Secchi |
|  | 99.15 | 90.65 | 92.92 | 92.39 | 86.92 | 91.30 | Secchi +<br>1 m |

Further examination of the specific Synechococcales ASVs revealed (1) seasonal patterns in ASV detection during ice-free periods, and (2) predominance of individual ASVs in the cyanobacterial community (e.g., ASV877 composed 57.7% of the surface community in July 2018, Big Turkey Lake; Figure 6). Across lake sites, eleven 16S rRNA gene ASVs were found to frequently comprise >10% of the cyanobacterial community alongside numerous low abundance ASVs. Notably, certain Synechococcales ASVs were found only seasonally, but others were ubiquitous across seasons. Examples of seasonally unique ASVs include: ASV886 ASV865 and ASV865 detected in higher abundances in seasonal transitory months of October and May, and ASV880 and ASV876 detected in higher abundances during summer sampling periods. Other ASVs such as ASV855 and ASV858 demonstrated seasonal ubiquity with detection across all sampling months including periods of ice-cover. The identification of seasonal and transition period specific ASVs indicates that a unique ecological niche was established during periods of lower water temperatures. Similar population diversity and niche development is evidenced with sequence variants that peaked during warmer summer temperatures.

**Figure 6:**
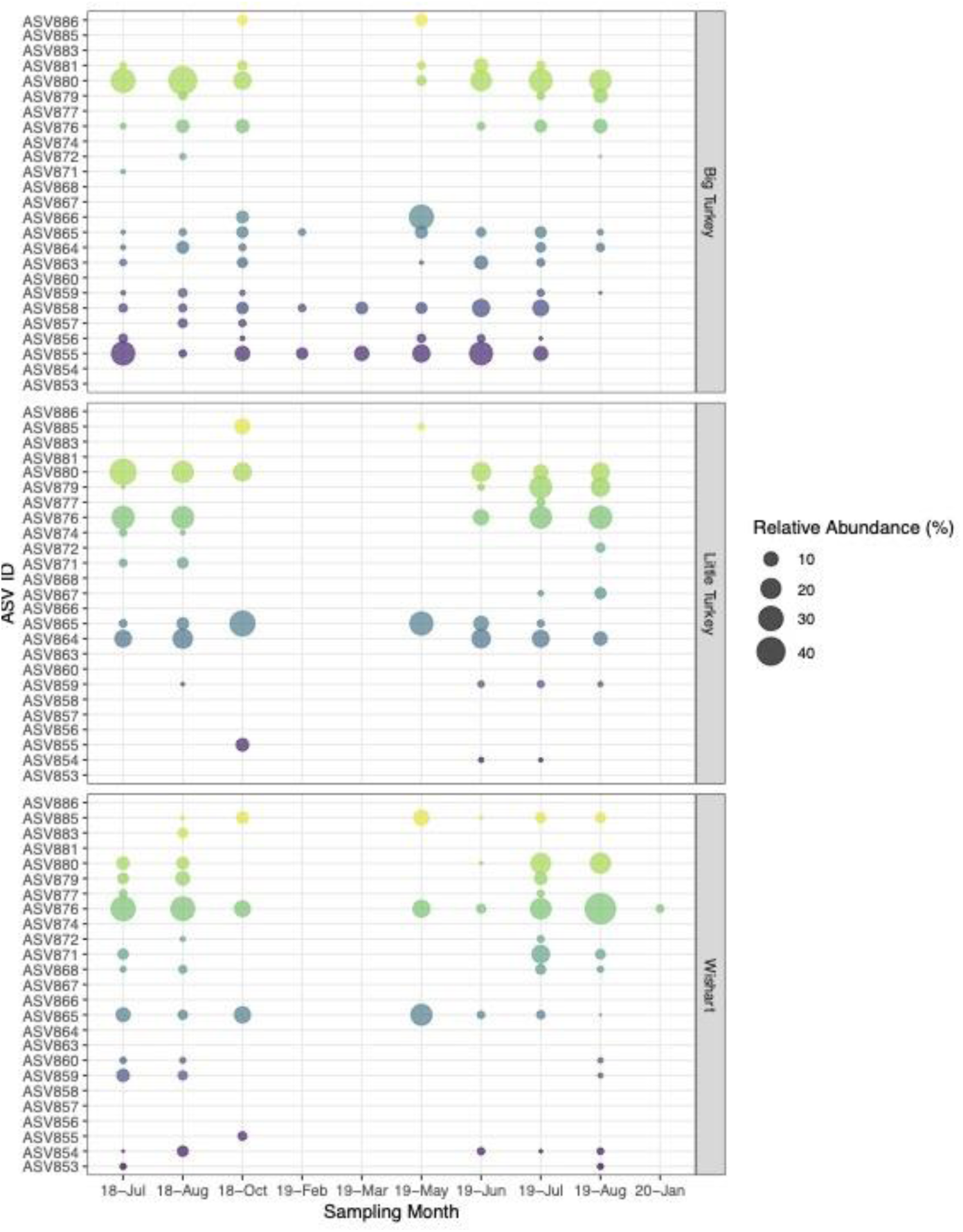
Contribution of individual amplicon sequence variants (ASVs) classified to the cyanobacterial order, Synechococcales across a long-term seasonal sampling series. Amplicon sequence variants present in less than 5% relative abundance cumulatively across all sampling months were excluded from visualization. Plotted points are scaled to represent relative abundance of individual ASVs in the cyanobacterial community demonstrating the seasonal occurrence and cycling of ASVs within the cyanobacterial community.

A range of cyanobacterial ASV distributions was also observed across sampling depths: some ASVs were detected exclusively at one depth, while others were distributed throughout the water column (Figure 7). Stratification affected the spatial distribution of ASVs, as would be expected. In Little Turkey and Big Turkey Lakes, the relative abundances of various ASVs were more heterogeneous than in Wishart Lake, which is shallow and does not stratify. This contrast is evident as the relative abundances of ASVs detected in Wishart are largely comparable across sampling depths. In the case of Big Turkey Lake, a large, stratified lake, ASV855 (July 2018) and ASV864 were detected in higher abundances at deeper sampling depths. Spatial variability in ASV distribution was also observed in Little Turkey Lake with the detection of ASV877, ASV867 and ASV860 in surface samples and the increasing relative abundance of ASV876 as sampling depth is increased.

**Figure 7:**
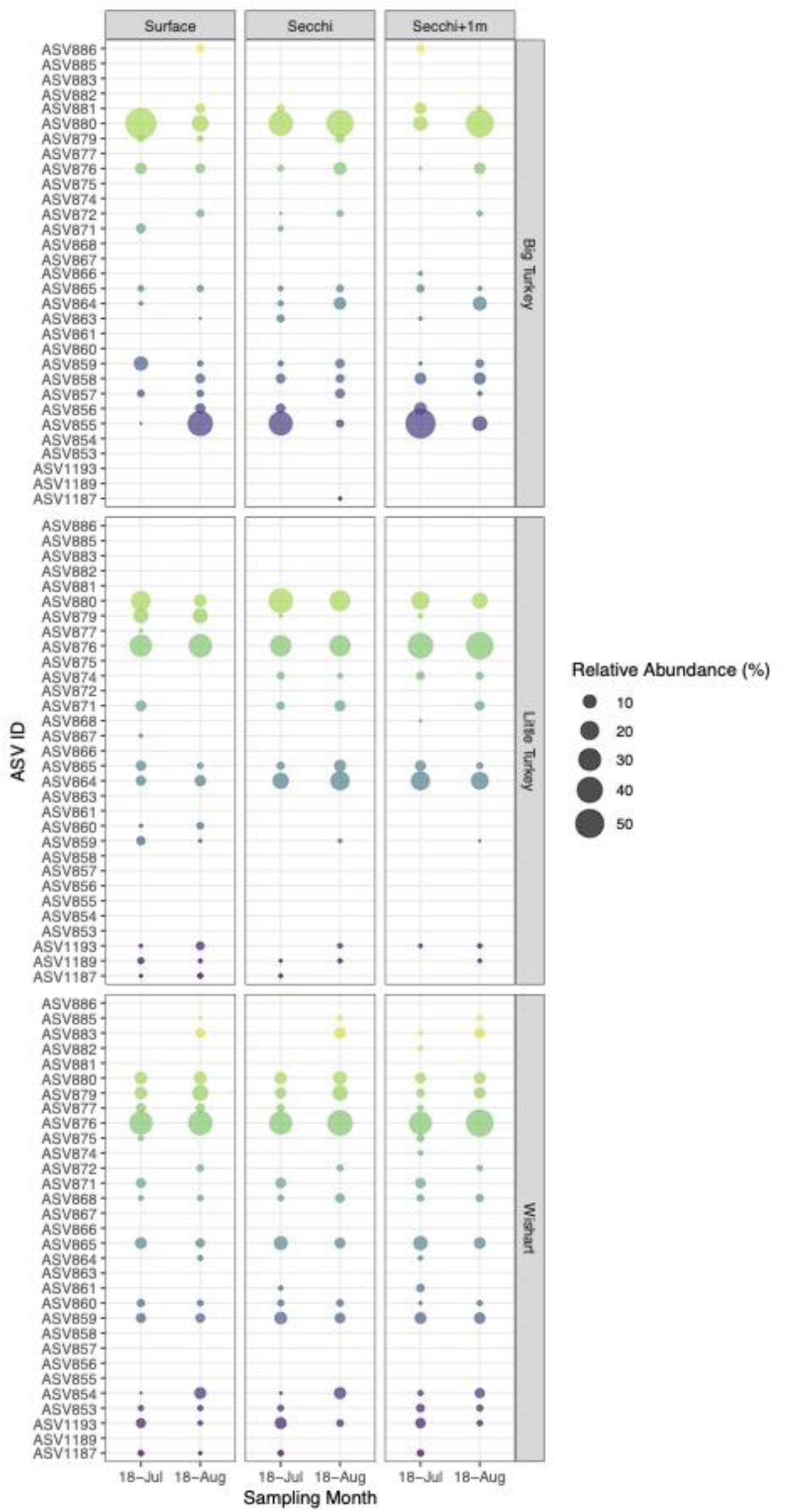
Contribution of individual amplicon sequence variants (ASVs) classified to the cyanobacterial order, Synechococcales across a water column profile. Variants present at >10% abundance in the cyanobacterial community are labelled to demonstrate individual dominance of cyanobacterial variants within the community. Plotted points are scaled to represent relative abundance of individual ASVs in the cyanobacterial community demonstrating the occupancy of spatial niches by certain ASVs.

### 3.4 Spatiotemporal Distribution of Cyanobacteria is System Specific

Bacterial community characterization across a water column depth profile in July and August 2018 demonstrated differences in cyanobacterial abundances (3.8 to 45.4%) between lakes (Figure 8; Table 2). Cyanobacterial sequences were significantly represented in the bacterial community across all sampling depths in Big Turkey and Wishart Lakes in July 2018 (*p* < 2.00e-05; Table S5; Figure S7). Specifically, within Big Turkey Lake, the highest cyanobacterial relative abundances were observed at the deepest sampling location (45.5%) indicating heterogeneous distribution across depths. Heterogenous distribution in the water column was also observed in Little Turkey Lake with cyanobacterial sequences only comprising a significant portion of the bacterial community at Secchi depth in July 2018 (*p* = 5.44e-05). In contrast, the relative abundance of cyanobacterial sequences was relatively homogenous (15.9 to 18.5% in July; 3.8 to 4.8% in August) across the shallow, non-stratified water column in Wishart Lake. Despite cyanobacteria composing a significant portion of the bacterial community in July 2018, by August 2018 cyanobacteria were not significantly represented across sampling depths in Wishart (*p* = 1), Little Turkey (*p* > 0.44) and Big Turkey (0.22 < *p* < 0.45) Lakes corresponding to a seasonal decrease in cyanobacterial populations within the bacterial community.

**Figure 8:**
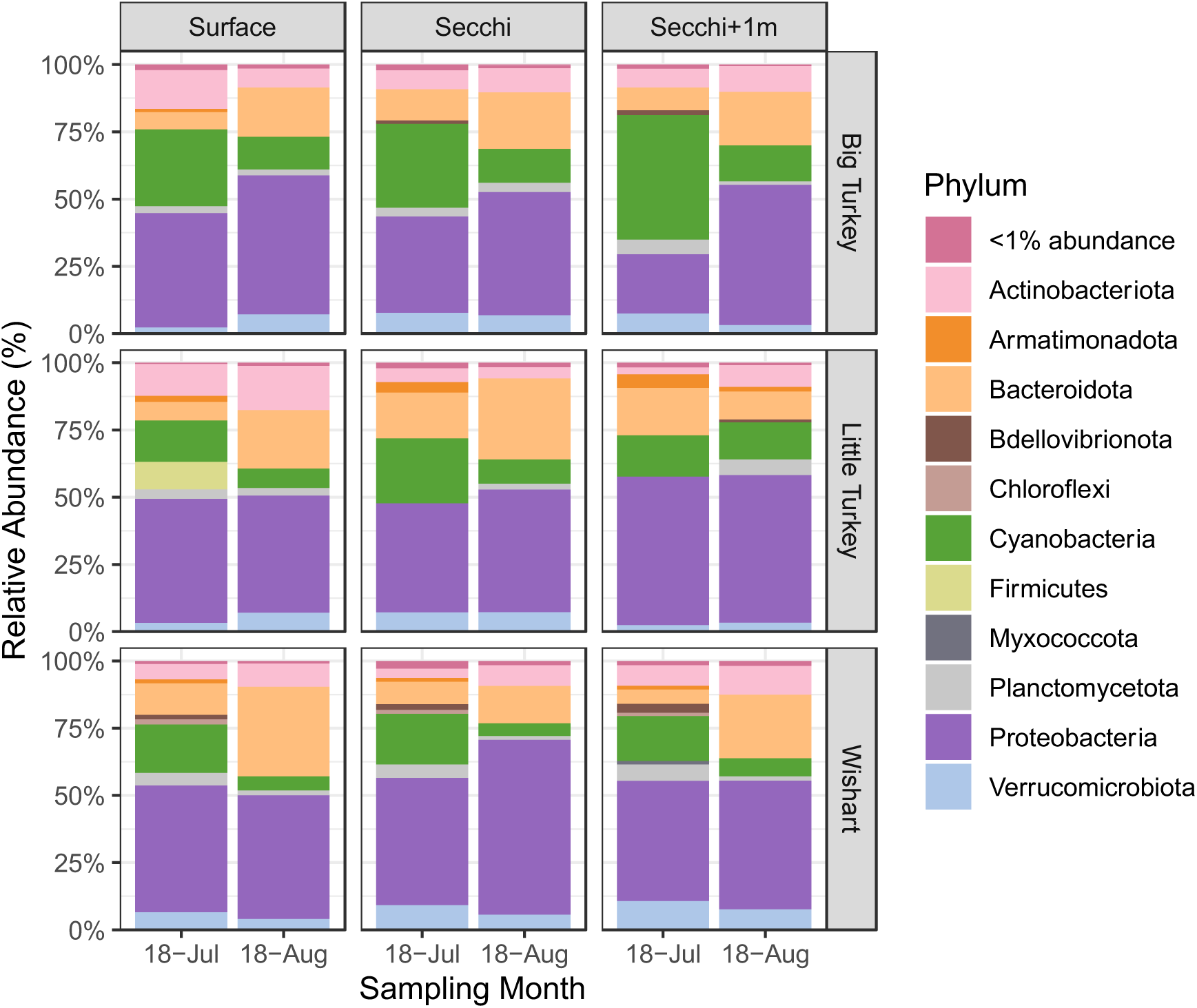
Stacked bar charts depicting the relative abundances of major bacterial phyla identified from amplicon sequencing of the V4 region of the 16S rRNA gene across a depth profile during the summer of 2018 in a non-stratified lake (Wishart), a mid-sized stratified lake (Little Turkey) and deep stratified lake (Big Turkey).

Differences in cyanobacterial community diversity across sampling depths and between summer sampling events in the three study lakes were also observed (Figure 9A). No spatial trends on diversity were distinguishable with variability seen in the sampling conditions producing the highest calculated Shannon Index. In Little Turkey Lake, Shannon Index was highest at the surface between both sampling months. Diversity across the water column of Wishart was comparable at all depths further supporting the potential for homogeneous cyanobacterial population distributions expected of a shallow, non-stratified lake. Distinct compositional similarities were also observed within lakes which may be influenced by stratification status. Across the shallow, well-mixed water column of Wishart Lake, cyanobacterial communities showed high similarity between sampling months, regardless of sampling depth (Figure 10). Little Turkey Lake also showed compositional similarity across sampling depths and sampling months. Big Turkey Lake community composition was the least similar, suggesting relatively greater variability in cyanobacterial community composition.

**Figure 9:**
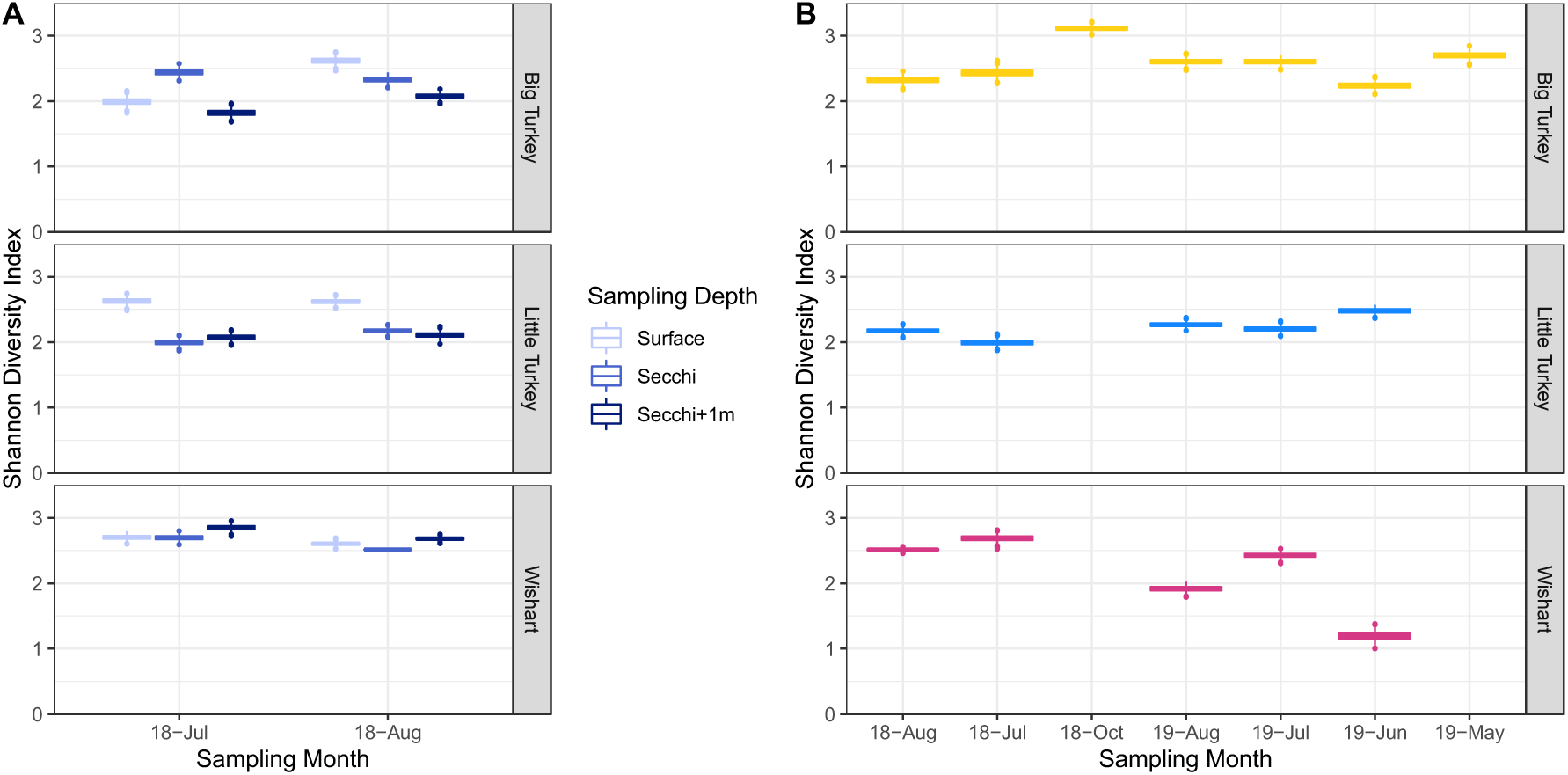
Alpha diversity analyses of cyanobacterial communities sampled during ice-free periods (A) across a water column depth profile in the summer of 2018, and (B) across a long-term seasonal timeline beginning in July 2018 and concluding in July 2019. Amplicon sequence variants classified to the phylum Cyanobacteria were selected to characterize diversity within cyanobacterial communities. The Shannon Index was calculated on rarefied library to evaluate the effects of (A) sampling depth and (B) sampling months on cyanobacterial communities between lake sites within the same watershed. Sampling months (B) that do not have data displayed represent instances where cyanobacterial sequences were present in low abundances and were excluded from diversity analyses.

**Figure 10:**
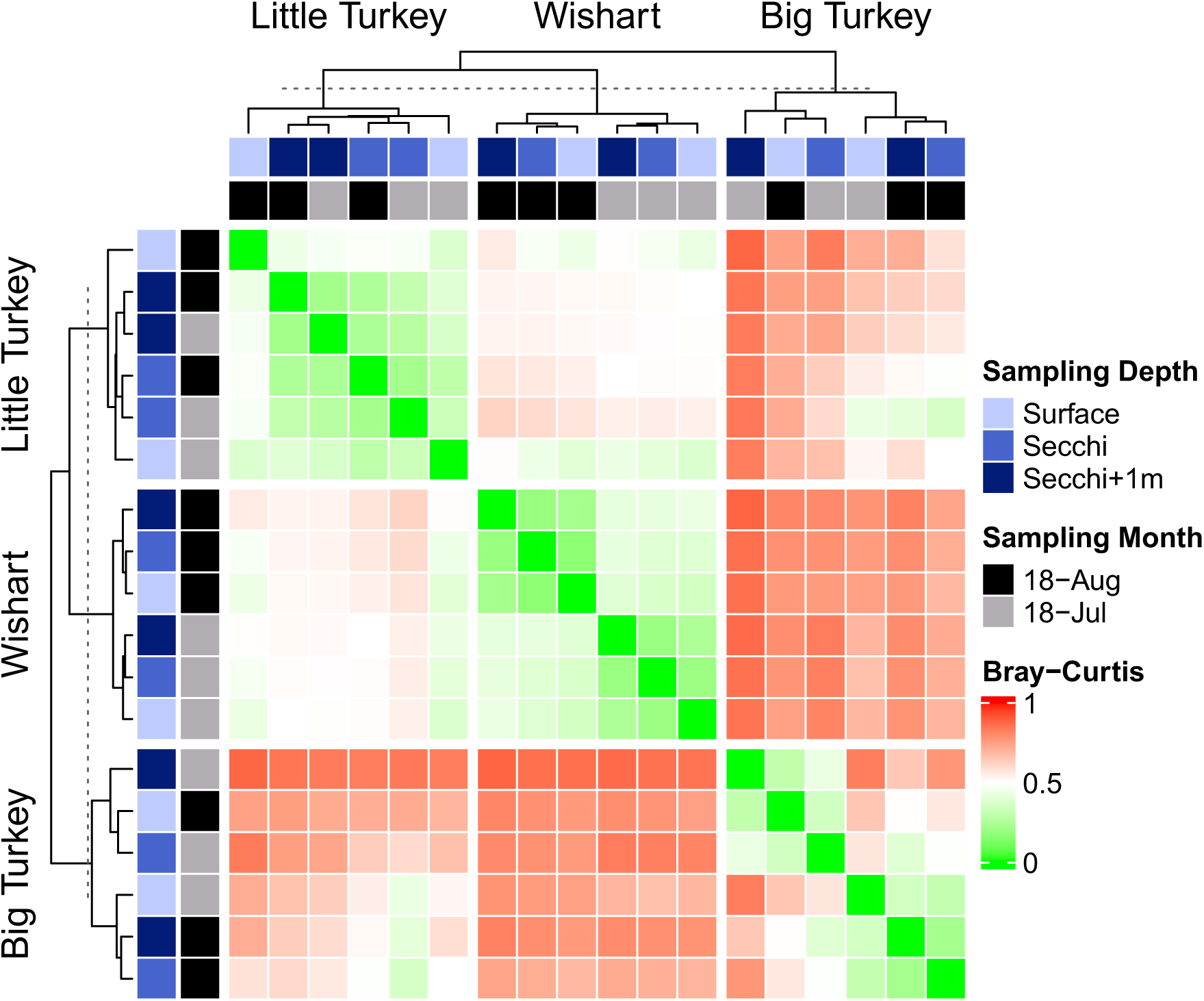
Compositional similarity analyses of cyanobacterial communities collected across the ice-free seasonal sampling series in a deep stratified (Big Turkey), mid-sized stratified (Little Turkey) and shallow non-stratified lake (Wishart). Bray-Curtis dissimilarity metric (0 = completely identical, 1 = completely dissimilar) was used to characterize similarity of cyanobacterial community sequence composition. Samples were collected across a water column depth (surface, Secchi, 1 metre below Secchi) in July and August 2018 to characterize trends in spatial variability between lakes of different thermal regimes.

During the period of lower productivity (i.e., October to May), the ability to examine seasonal trends in diversity was limited by low sequence counts. Nonetheless, seasonal trends in community diversity were observed between ice-free sampling months. In all lakes, Shannon Index values did not show major fluctuations throughout the ice-free sampling period (Figure 9B). The full ice-free sampling period in Big Turkey Lake enabled identification of potential transitions in cyanobacterial community composition in response to seasonal shifts (Figure 11). Cyanobacterial communities in May 2019 from Big Turkey Lake were most dissimilar from other sampling months within and across lakes which may represent transitionary seasonal shifts in community composition. Comparatively, summer months (June – August) into October 2019 were more similar within Big Turkey Lake. The occurrence of seasonally specific community composition is also observed in Wishart and Little Turkey Lake with high similarity observed within both lakes between sampling years (2018 – 2019) in July and August.

**Figure 11:**
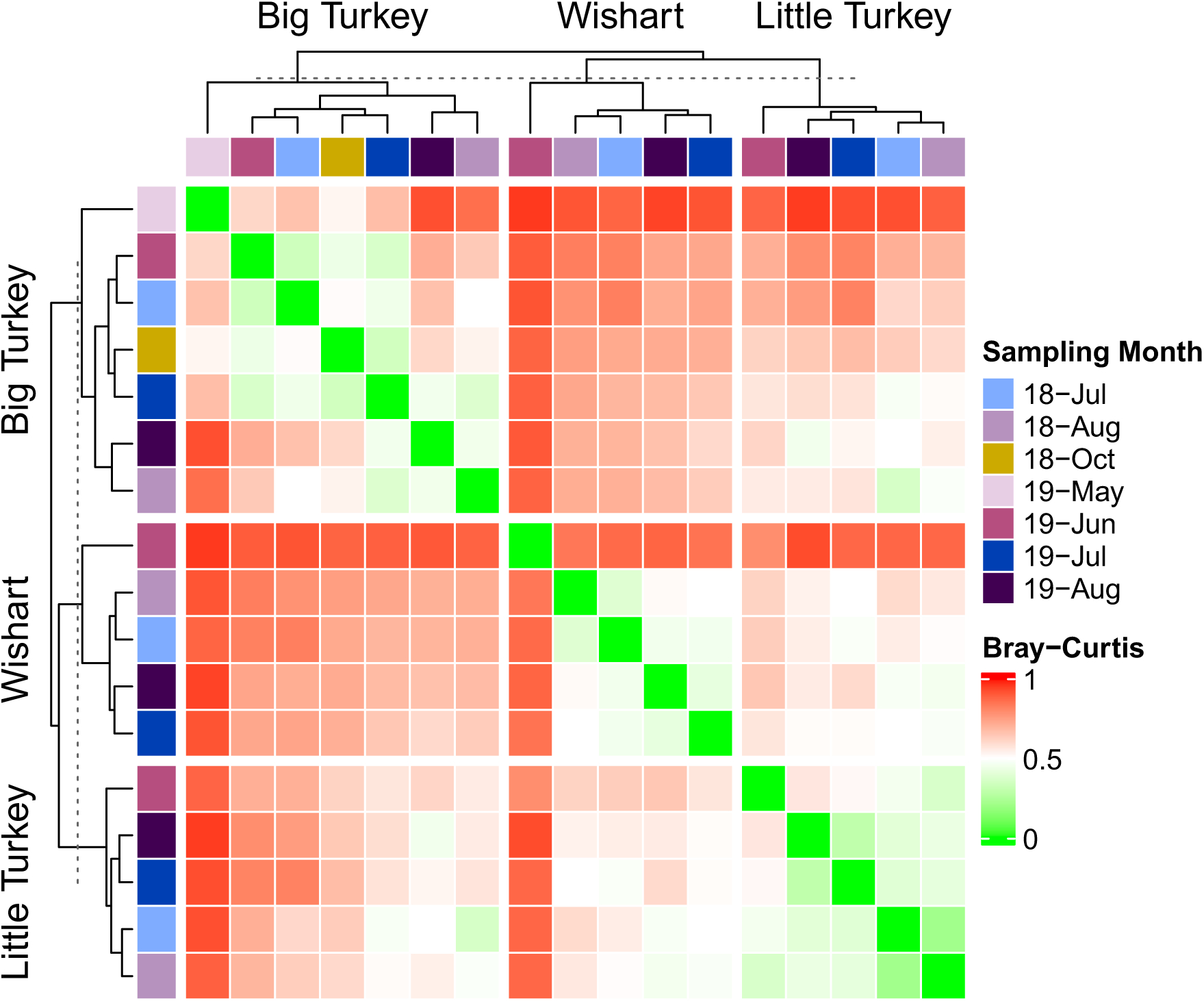
Compositional similarity analyses of cyanobacterial communities collected across the ice-free seasonal sampling series in a deep stratified (Big Turkey), mid-sized stratified (Little Turkey) and shallow non-stratified lake (Wishart) across a long-term seasonal profile. Bray-Curtis dissimilarity metric (0 = completely identical, 1 = completely dissimilar) was used to characterize similarity of cyanobacterial community sequence composition. Samples were collected monthly during ice-free periods commencing in July 2018 and concluding in August 2019 to characterize annual seasonal shifts in cyanobacterial community composition.

## 4. Discussion, Conclusions & Implications

Consistent with global trends (Carey et al., 2008; Wells et al., 2020) the number of algal blooms reported in Ontario, Canada has been significantly increasing, especially in lakes on the Canadian Shield where these increases are predominantly composed of potentially toxin-producing cyanobacteria (Winter et al., 2011). The multi-year bacterial community analysis conducted across three lakes in the TLW and reported herein aligns with those observations. Lingering beliefs regarding winter limnology often dismiss winter periods (especially ice cover) as ecologically unimportant relative to the summer growing season (Powers & Hampton, 2016). Contrasting this belief, we demonstrate that evaluation of broader seasonal variation in lake microbial communities can provide critical insights regarding climate change impacts on oligotrophic, northern temperate lake ecosystems. This may include identification of the associated implications to human and environmental health through the characterization of cyanobacterial community dynamics in a changing world. This study provided seven important observations:

(1) almost 40 years of ice phenology data showed that rising air temperatures have led to significantly longer ice-free periods extending later into the year annually in the TLW,
(2) warmer spring air temperatures, increased solar radiation and elevated discharge rates following snowmelt coincided with warming of the water column and initiated the aquatic growing season,
(3) cyanobacteria persisted year-round in the oligotrophic, northern temperate lakes of the TLW,
(4) cyanobacterial communities during ice-covered months included sequences classified to the recently identified non-photosynthetic, potentially toxic basal lineage, Melainabacteria,
(5) cyanobacteria composed a significant portion of the bacterial communities in the study lakes during ice free periods as early as May and persisted into late October,
(6) picocyanobacteria were especially dominant during ice-free periods,
(7) individual picocyanobacterial populations shifted seasonally—while certain sequences were dominant during ice-free months, other sequences were restricted to either (i) the shoulder seasons of the ice-free period (i.e., spring and fall) or (ii) periods of winter ice cover, and
(8) lakes with lower depth ratios and longer water renewal times situated in areas relatively rich in fine grained sediment (i.e., Big Turkey Lake; Table S2) had higher relative abundances of cyanobacteria.

Future increases in air temperature are expected to cause further decreases in days of lake ice cover (Woolway et al., 2022), a phenomenon which is being observed in the lakes of TLW. However, in the case of the lakes of TLW, this decrease is caused by a later ice-on date in the autumn where cyanobacteria were not observed in high relative abundances which contrasts with the observance of significant proportions of cyanobacteria as early as May in some systems. This suggests that potentially earlier loss of ice cover in the spring may impact photosynthetic growth more significantly than an extension of a longer aquatic growing season in the autumn. Winter periods for northern temperate lakes such as those used in this study would have significantly different environmental conditions not favouring photosynthetic growth. Specifically, solar radiation input would be drastically reduced due to the presence of thick ice cover and snow cover limiting photosynthetic activity (Bertilsson et al., 2013). However, the loss of snow cover, an event which will naturally occur with melting due to increased solar radiation and warmer temperatures in the early spring, will result in warming of surface waters and convective mixing (Bertilsson et al., 2013). While such phenomena are expected to occur in the lakes of TLW, greater resolution in data is required to fully characterize the underlying drivers impacting the seasonal transitions of the physicochemical conditions in these lakes. With available data and based on previous research on seasonality in lakes, it is expected that the combination of physical (*i.e.* temperature increase, increased solar radiation) and chemical (*i.e.* access to nutrients through convective mixing) changes gives cyanobacteria a competitive advantage to thrive in seasonal shifting environments observed during the spring due to their ability to grow in low-temperature, low-light and low nutrient environments (Reinl et al., 2021).

The presence and persistence of potentially toxin forming picocyanobacterial taxa within lakes of the temperate forest biome of Canada have not previously been reported. In the lakes of TLW, picocyanobacterial taxa dominated cyanobacterial communities seasonally (Figure 4) and spatially (Figure 5), often comprising a statistically significant portion of the bacterial community during ice-free periods. This demonstrates their exploitation of environmental variability through evolutionary mechanistic adaptations (e.g., small size, ability to grow in low light intensity environments, rapid nutrient uptake, ability to maintain water column position, etc.) and interactions (e.g., allelopathy) with larger primary producers and predators. Although picocyanobacteria are abundant in diverse freshwater and marine environments (Cai et al., 2010; Chen et al., 2006; Collos et al., 2009; Felföldi et al., 2016; Gin et al., 2021) sufficient understanding of the occurrence and characterization of their blooms, toxicity, and allelopathic activity is lacking (Śliwińska-Wilczewska et al., 2018), especially in freshwater environments. Abiotic and biotic factors such as lake morphometry, thermal regime, and trophic state influence picocyanobacterial dynamics and lakes in close proximity may exhibit varying seasonal population trends (Ruber et al., 2018). Specifically, lakes with lower depth ratios and longer water renewal times situated in areas rich in fine sediments that have the potential to deliver limiting nutrients (i.e., phosphorus) are especially vulnerable to increased primary productivity, including the proliferation of potentially harmful algae. It has been suggested that these factors may be at least as important as nutrients in affecting the structure of freshwater picocyanobaterial communities (Callieri, 2008).

Picocyanobacteria are common components of the photic zone, but distribution and abundance can range widely depending on the system conditions ((Callieri & Stockner, 2002). Previous reports including a survey of 43 lakes and ponds indicated that picocyanobacteria prefer large, deep lakes with high hydrologic retention times and incomplete mixing due to vertical density differences (Callieri, 2008; Camacho et al., 2003). The present investigation was consistent with such observations—in TLW, higher relative abundances of cyanobacteria were detected in the stratified lakes (Big Turkey and Little Turkey Lakes) compared to the shallow, well-mixed lake (Wishart Lake) (Figure 3; Table 1). Picocyanobacteria have previously been observed to reach peak abundances prior to the onset of thermal stratification, (Callieri & Stockner, 2000; Fahnenstiel et al., 1991; Li et al., 2020), but herein Synechococcales sequences were found to be predominant in lakes year-round independent of the onset of thermal stratification. Highly variable vertical distribution of populations within thermally stratified lakes has previously been reported (Hall & Vincent, 1994; Stockner et al., 2006) and samples collected at different depths within the metalimnion of Big Turkey and Little Turkey Lake also showed variability in picocyanobacterial 16S rRNA gene ASVs suggesting heterogeneity in water column distribution. In much larger lakes such as Lakes Huron and Michigan, peak abundances of picocyanobacteria have been detected in the lower metalimnion and upper hypolimnion (Stockner et al., 2006) with higher abundances occurring at low light intensity (Jakubowska & Szeląg-Wasielewska, 2015; Pick & Agbeti, 1991). Consistent with these reports of occurrence in the lower metalimnion with low light intensity, the highest proportion of cyanobacterial sequences within the bacterial community in the present investigation was in Big Turkey Lake one metre below Secchi depth (Figure 8; Table 2).

In temperate freshwater and marine environments, picocyanobacteria are typically more abundant in the warm season than in the cold season, during which cell density decreases of approximately three orders of magnitude (Postius & Ernst, 1999; Waterbury & Valois, 1993) and shifts to completely different populations (Cai et al., 2010b) have been reported. Similar bimodal patterns and shifts between summer and winter relative abundance of picocyanobacteria were observed in the TLW, as reported herein. These seasonal shifts in populations were observed in the ASV composition of cyanobacterial communities (Figure 6; Figure 7) due to specific populations or subclades being more adapted to lower temperatures (Cai et al., 2010) resulting in the non-ubiquitous occurrence of ASVs.

The significant representation of picocyanobacteria in the bacterial community of Big Turkey Lake shortly after the spring snowmelt in May 2019 is especially notable. The warming climate has led to increased air temperatures in TLW (Webster et al., 2021). This, coupled with the relatively greater availability of fine-grained sediment (i.e., glacial till) surrounding the lower elevation lakes of the watershed, the shifts in precipitation patterns, and the specific lake morphometry of Big Turkey Lake as described by Jeffries et al. (1988) may explain the significant spring proliferation (Carpenter, 1983; Genkai-Kato & Carpenter, 2005). The depth ratio (i.e., the ratio of mean to maximum lake depth) is substantially lower in Big Turkey Lake than in the other study lakes. In lakes such as Big Turkey Lake (Figure S2; Table S1), in which the thermocline is shallower than approximately one to two times the mean depth, the epilimnion’s sediment surface area to volume ratio declines with depth ratio. The potential nutrient recycling from the sediment surface, productivity, and sediment accretion rates are expected to increase as depth ratio decreases (Carpenter, 1983) suggesting higher nutrient recycling present in Big Turkey Lake relative to the other lakes in the watershed. As well, more sediment can be eroded during runoff from the Big Turkey Lake watershed during precipitation events or snowmelt periods because there is more available sediment on the surrounding landscape, relative to Wishart Lake (Jeffries et al., 1988). The delivery of available sediment to oligotrophic lakes such as Big Turkey Lake may also result in the release of phosphorus to the water column (Withers & Jarvie, 2008) and contribute to the spring proliferation of picocyanobacteria (Passoni & Callieri, 2000) due to their efficient nutrient utilization (Śliwińska-Wilczewska et al., 2018). Nutrient availability in the lake is further supported by the abundance of macrophytes at its margins (Jefferies et al., 1988), which may further modulate phosphorus recycling from sediments (Genkai-Kato and Carpenter, 2005). Changes in the biotic or abiotic factors that would alter this complex balance at the watershed-scale cannot be described at present and warrant further investigation.

The presence and dominance of picocyanobacteria in TLW is further notable due to their ability to produce several metabolites of human and environmental health concern or aesthetic significance. They can produce the hepatotoxins microcystin and nodularin (Chorus et al., 2000; Jakubowska & Szeląg-Wasielewska, 2015; Vareli et al., 2013). Several species of picocyanobacterial may also synthesize the neurotoxin β-N-methylamino-L-alanine (BMAA) (Cervantes Cianca et al., 2012; Cox et al., 2003), which is implicated in neurodegenerative diseases such as Parkinson’s and Alzheimer’s (Cervantes Cianca et al., 2012). They can also produce cylindrospermopsin and anatoxin with toxicity often underestimated due to assay sensitivity limitations (Gin et al., 2021). In addition to toxins, picocyanobacteria may also produce geosmin and MIB (Graham et al., 2008; Jakubowska & Szeląg-Wasielewska, 2015; Watson, 2003), which are taste and odour forming compounds that commonly result in customer complaints when present in distributed drinking water (McGuire, 1995; Suffet et al., 1996). While picocyanobacteria significantly contribute to total primary productivity in freshwater and marine environments (Stockner et al., 2002; Waterbury et al., 1986), the biotic and abiotic factors that drive their proliferation and potential toxin production are not well understood (Śliwińska-Wilczewska et al., 2018).

In addition to picocyanobacteria, Melainabacteria and Sericytochromatia sequences were also found at TLW. They have been previously observed in aphotic environments (Monchamp et al., 2019a; Soo et al., 2014). Although their biogeography and ecology remain understudied, genomes have previously been isolated from various waters including lakes (Monchamp et al., 2019a) and engineered aquatic systems including water treatment facilities and water distribution systems (Ling et al., 2018; Zamyadi et al., 2019). The presence of Melainabacteria and Sericytochromatia sequences in TLW when lakes were covered with up to 0.81 m of ice and additional snow coverage limiting light availability aligns with those previous reports. The detection of this non-phototrophic basal lineage of Cyanobacteria during periods of ice cover suggests their potential role in winter microbial community processes. While the ecological function and role of these organisms is not well understood (Monchamp et al., 2019b), synthesis of BMAA has been speculated (Nunes-Costa et al., 2020), emphasizing the need for further investigation of their distribution and function.

Given the human and environmental health implications of these compounds and the significant costs associated with their removal from drinking water (Emelko et al., 2011), it is critical to explore the possibility of picocyanobacteria-associated toxin production in freshwaters. This production may be undetected because of reliance on visual observation of accumulated cyanobacterial biomass and traditional foci on monitoring of colonial and filamentous bloom forming cyanobacteria (Chorus et al., 2000; Water Quality Research Australia & Global Water Research Coalition, 2009). Both cyanobacterial cell accumulation and toxin production have been observed at water treatment plants in absence of visible blooms in source water supplies (Almuhtaram et al., 2018) underscoring that picocyanobacterial and other non-traditional cyanotoxin producers (e.g., Melainabacteria, Sericytochromatia) may be unnoticed (because water providers are not required to test for toxin production in absence of visible blooms) and thus, under-reported. Accordingly, broader and more comprehensive monitoring is required (Pobel et al., 2011; Welker et al., 2021) to advance understanding of picocyanobacterial and non-photosynthetic cyanobacteria population dynamics in response to local biotic and abiotic drivers, some of which are impacted by changing climate (Callieri, 2008; Drakare & Liess, 2010).

While climate change is not being proposed herein as a driver of the persistence and dominance of picocyanobacteria in TLW, it emphasizes the pressing need to better understand the potential roles that picocyanobacteria and non-photosynthetic Melainabacteria (such as those associated with sequences observed in TLW) may play in toxin production and trophic status modulation, especially in oligotrophic lakes and reservoirs that are relied upon for the provision of drinking water. It is generally understood that algal blooms occur at the height of summer and in early fall. However, the early re-emergence of cyanobacteria in the microbial community in the spring despite non-significant shifts in the start of the ice-off period suggests the critical importance of snowmelt and the warmer post-winter temperatures to initiating the aquatic growing season and allowing for proliferation of cyanobacterial populations in northern, temperate, oligotrophic lakes. This persistence may be expected to continue into the autumn as a result of additional nutrient loading from autumn storms shifting to the post-canopy leaf fall period in these biomes (Creed et al., 2015). Although significantly higher precipitation has been observed in the autumn, cyanobacterial populations did not exhibit the same microbial community dominance as that observed in the spring suggesting that the lower temperatures and lower discharge rates in the autumn cannot support the same levels of cyanobacterial growth as that of the spring. Notably, photosynthetic populations of potentially toxic cyanobacteria reappeared in the water column immediately following the loss of ice cover. They were especially abundant in lakes where surficial geology and lake morphometry favor greater availability of fine sediment and associated nutrients further highlights the impact of snowmelts on oligotrophic lakes primary productivity. Thus, this collective analysis demonstrates that the convergence of key abiotic and biotic factors— climate forcing of hydrological and biogeochemical processes, and intrinsic landscape features—may enable increases in the relative abundance of potentially toxic cyanobacteria (i.e., picocyanobacteria) within the temperate forest biome of Canada over increasingly longer periods of time. More detailed, year-round monitoring is needed to better characterize climate change-exacerbated shifts in algal community dynamics and manage associated public health and aquatic ecosystem risks.

## 5. Acknowledgments

We acknowledge the support of NSERC (Natural Science and Engineering Research Council of Canada), specifically through the *for*Water NSERC Network for Forested Drinking Water Source Protection Technologies (NETGP-494312-16). This research was undertaken, in part, thanks to funding from the Canada Research Chairs Program (ME; Canada Research Chair in Water Science, Technology & Policy). We are grateful for the support from Natural Resources Canada (NRCAN), the Great Lakes Forestry Centre (Canadian Forest Service) and Environment and Climate Change Canada (ECCC) in sample collection at the Turkey Lakes Watershed. We would also like to acknowledge Roy Neureuther’s (ECCC, retired) dedication to observing and recording lake ice conditions at the Turkey Lakes Watershed during his career.

