## Supplementary Figures & Tables for "Early Seasonal Increases and Persistence in Relative Abundance of Potentially Toxic Cyanobacteria: Concerning Impacts of Extended Ice-Free Periods in Northern Temperate Lakes"

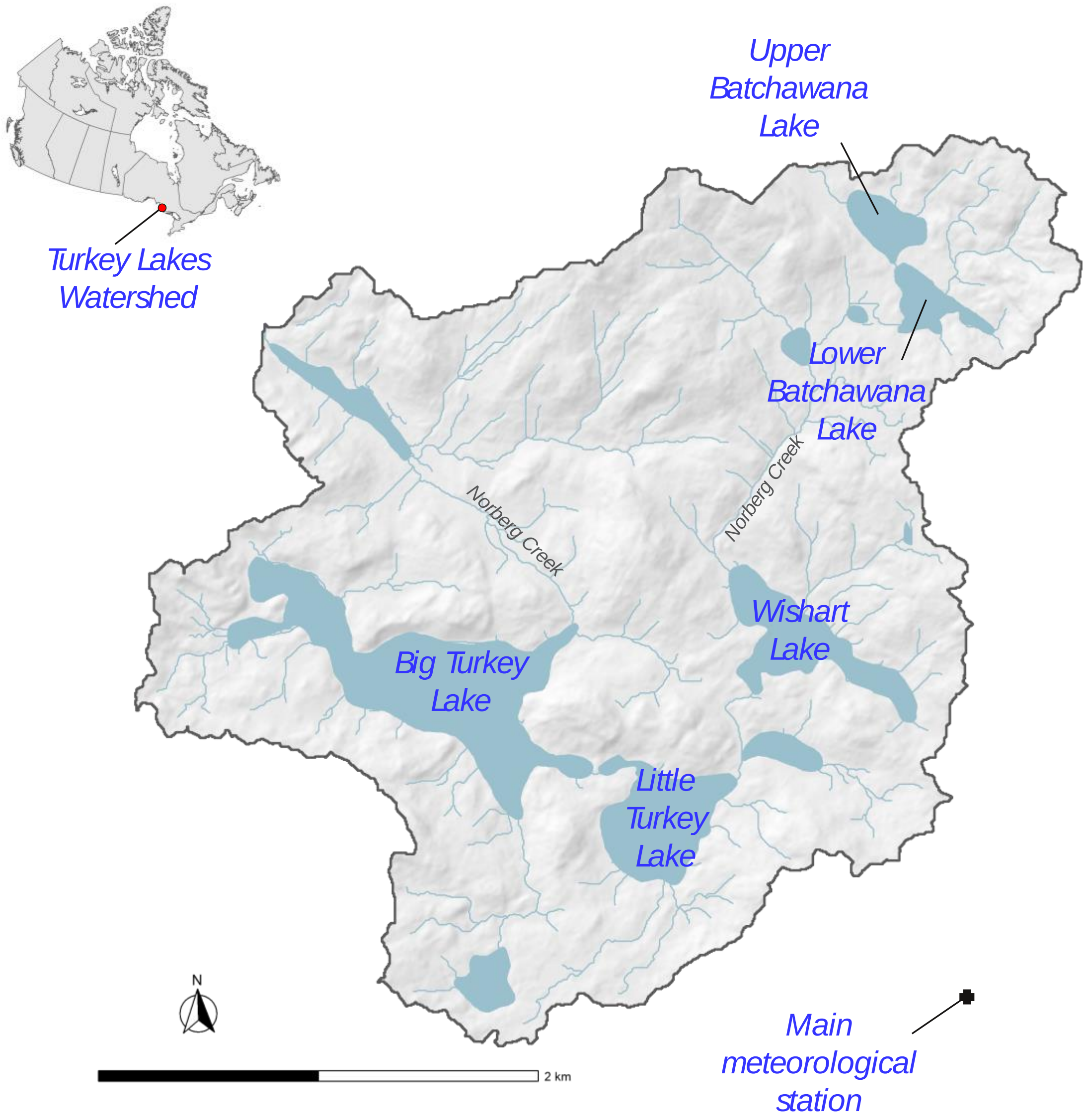

**Figure S1** The Turkey Lakes Watershed location and lake sites. The watershed is located approximately 50 km north of Sault Ste Marie, Ontario as indicated by the marker on the map.

**Table S1** Summary of characteristics of the lakes of Turkey Lakes Watershed adapted from Jeffries et al. (1988).

| Lake | Drainage Basin Area (ha) | Lake Surface Area (ha) | Maximum Depth (m) | Mean Depth (m) | Lake Volume ($m^3 * 10^5$) | Water Renewal Time (yr) |
|---|---|---|---|---|---|---|
| Upper Batchwana | 24.0 | 5.88 | 11.3 | 3.87 | 2.27 | 1.3 |
| Lower Batchwana | 85.6 | 5.82 | 10.9 | 3.27 | 1.90 | 0.30 |
| Wishart | 337 | 19.2 | 4.5 | 2.19 | 4.21 | 0.15 |
| Little Turkey | 491 | 19.2 | 13.0 | 6.04 | 11.6 | 0.25 |
| Big Turkey | 803 | 52.0 | 37.0 | 12.2 | 63.4 | 0.94 |

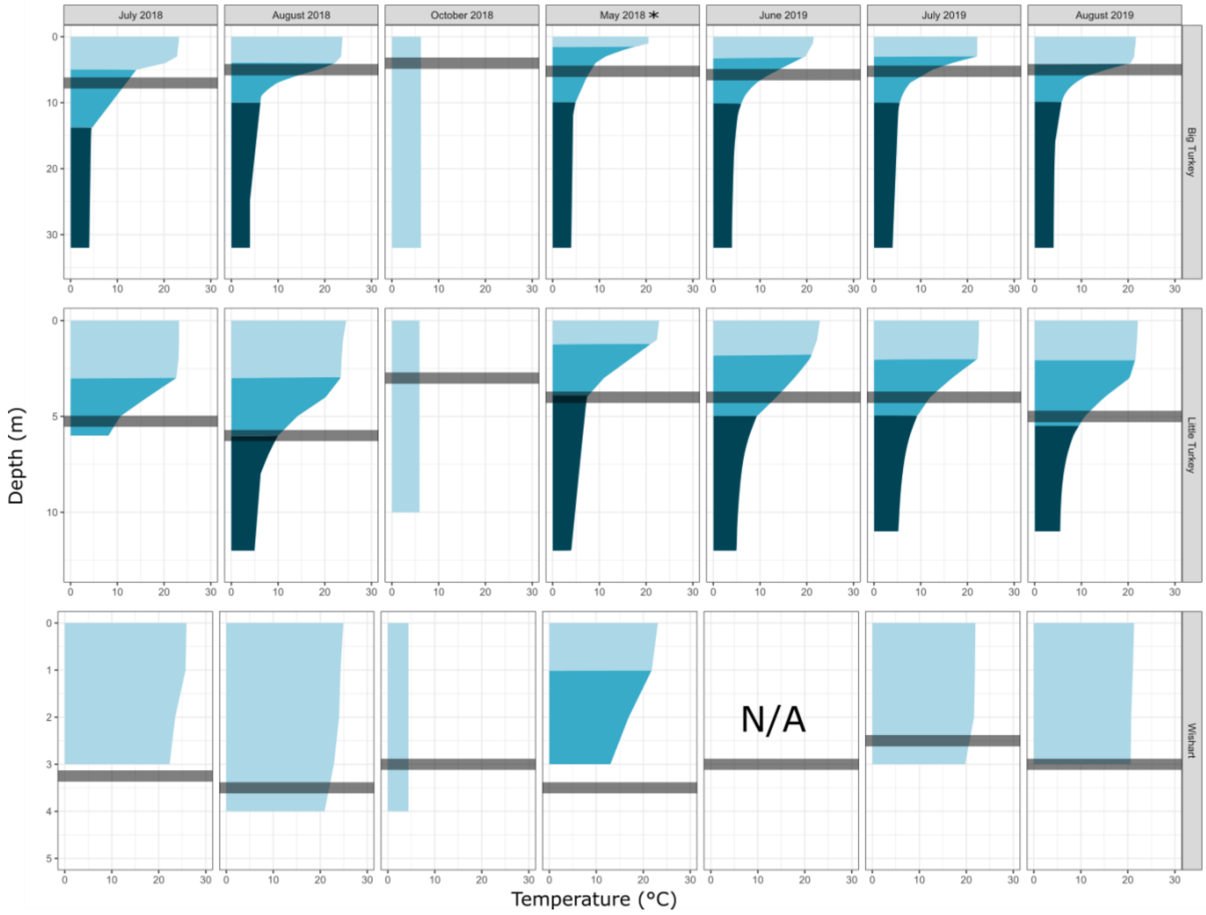

**Figure S2** Water column temperature profiles in ice-free months. Water column temperature profiles were collected during ice-months in Big Turkey (Max depth = 37 m), Little Turkey (Max depth = 13 m) and Wishart Lake (Max depth = 4.5 m). Secchi depth, which is used as a sampling depth in the studies in the seasonal profiling of this research, is indicated with the shaded grey bar. Thermally stratified layers are identified as epilimnion (light shade), metalimnion (medium shade) and hypolimnion (dark shade). Notably, water column profiles for June 2019 in Wishart Lake were not available. May 2019 data in all three lake sites were also unavailable but data from May 2018 have been provided to demonstrate previous thermal stratification trends.

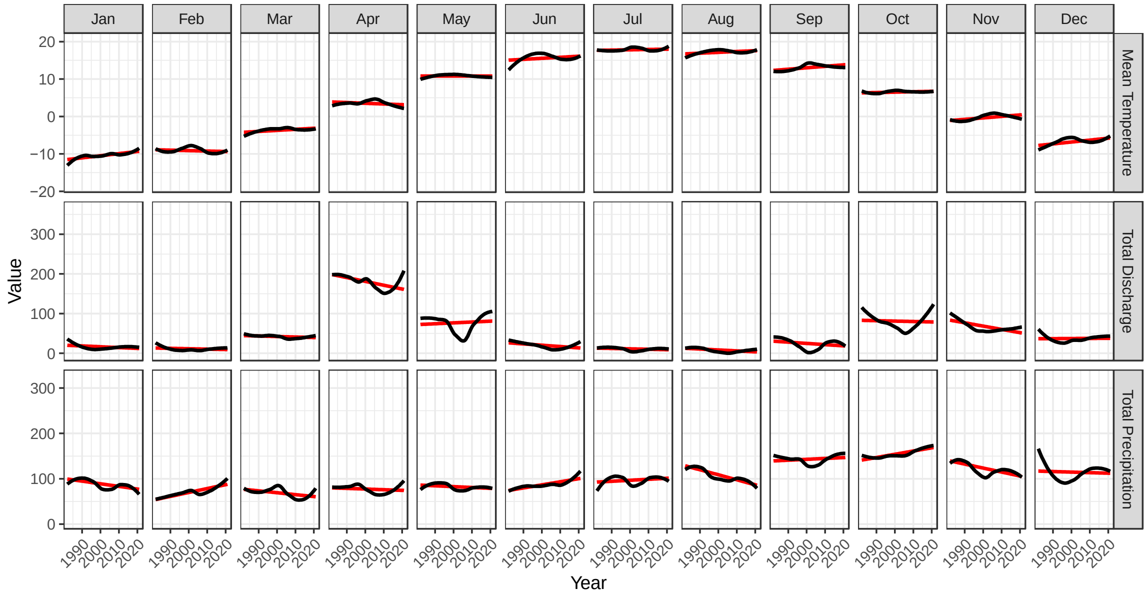

**Figure S3:** Hydroclimatic trends in the Turkey Lakes Watershed measured from 1982-2021.

**Table S2** Sample summary for seasonality and depth profile for bacterial community characterization. All water samples were collected using a peristaltic pump (Masterflex E/S Portable Sampler), then vacuum filtered through a 47 mm GF/C filter (Whatman, plc, Buckinghamshire, United Kingdom), and stored at -20°C prior to DNA extraction. To ensure sufficient biomass for analysis, 350 to 1000 mL of water were filtered.

| Sample ID | Lake | Sampling Depth (m) | Sampling Date | Volume Filtered (mL) | DNA Concentration (ng/μL) |
|---|---|---|---|---|---|
| TLW37 | Big Turkey | 0 | July 18, 2018 | 950 | 2.2 |
| TLW42 | Big Turkey | 7 | July 18, 2018 | 800 | 12.4 |
| TLW43 | Big Turkey | 8 | July 18, 2018 | 800 | 11.7 |
| TLW82 | Big Turkey | 0 | August 13, 2018 | 750 | 8 |
| TLW85 | Big Turkey | 5 | August 13, 2018 | 600 | 12.1 |
| TLW88 | Big Turkey | 6 | August 13, 2018 | 500 | 8.7 |
| TLW171 | Big Turkey | 4 | October 25, 2018 | 1000 | 7.9 |
| TLW181 | Big Turkey | 0.85 (Under Ice) | February 19,2019 | 1000 | 10.3 |
| TLW191 | Big Turkey | 1.06 (Under Ice) | March 25, 2019 | 1000 | 12.7 |
| TLW232 | Big Turkey | 5.25 | May 23, 2019 | 1000 | 7.4 |
| TLW277 | Big Turkey | 5.75 | June 28, 2019 | 1000 | 0.9 |
| TLW322 | Big Turkey | 5.25 | July 24, 2019 | 1000 | 0.7 |
| TLW325 | Big Turkey | 6.25 | July 24, 2019 | 1000 | 1.6 |
| TLW367 | Big Turkey | 5 | August 21, 2019 | 1000 | 0.9 |
| TLW407 | Big Turkey | N/A (Under Ice) | January 23, 2020 | 1000 | 1.2 |
| TLW28 | Little Turkey | 0 | July 18, 2018 | 1000 | 3.3 |
| TLW31 | Little Turkey | 5.25 | July 18, 2018 | 400 | 2.8 |

| **Table S2 Continued** | | | | | |
|---|---|---|---|---|---|
| Sample ID | Lake | Sampling Depth (m) | Sampling Date | Volume Filtered (mL) | DNA Concentration (ng/µL) |
| TLW36 | Little Turkey | 6.25 | July 18, 2018 | 500 | 2.8 |
| TLW73 | Little Turkey | 0 | August 13, 2018 | 1000 | 4.1 |
| TLW76 | Little Turkey | 6 | August 13, 2018 | 550 | 5.7 |
| TLW79 | Little Turkey | 7 | August 13, 2018 | 600 | 4.7 |
| TLW169 | Little Turkey | 3 | October 25, 2018 | 1000 | 3.3 |
| TLW179 | Little Turkey | 0.89 (Under Ice) | February 19, 2019 | 1000 | 4.4 |
| TLW189 | Little Turkey | 1.03 (Under Ice) | March 25, 2019 | 1000 | 2.4 |
| TLW223 | Little Turkey | 4 | May 23, 2019 | 1000 | 6 |
| TLW268 | Little Turkey | 4 | June 28, 2019 | 1000 | 2.2 |
| TLW313 | Little Turkey | 4 | July 24, 2019 | 1000 | 1.4 |
| TLW358 | Little Turkey | 5 | August 21, 2019 | 1000 | 2.2 |
| TLW409 | Little Turkey | 0.76 (Under Ice) | January 22, 2020 | 1000 | 1.8 |
| TLW19 | Wishart | 0 | July 16, 2018 | 950 | 3 |
| TLW22 | Wishart | 3.25 | July 16, 2018 | 400 | 3.5 |
| TLW26 | Wishart | 4.25 | July 16, 2018 | 250 | 2.6 |
| TLW64 | Wishart | 0 | August 14, 2018 | 600 | 0.8 |
| TLW67 | Wishart | 3.5 | August 14, 2018 | 450 | 0* |
| TLW70 | Wishart | 4.5 | August 14, 2018 | 350 | 3.2 |
| TLW167 | Wishart | 3 | October 25, 2018 | 1000 | 2.5 |
| TLW177 | Wishart | 0.87 (Under Ice) | February 19, 2019 | 1000 | 2.6 |
| TLW187 | Wishart | 0.96 (Under Ice) | March 27, 2019 | 1000 | 5.5 |
| TLW214 | Wishart | 3.5 | May 22, 2019 | 1000 | 3.1 |
| TLW259 | Wishart | 3 | June 28, 2019 | 600 | 6.3 |
| TLW304 | Wishart | 2.5 | July 25, 2019 | 500 | 3.5 |

**Table S2 Continued**

| Sample ID | Lake | Sampling Depth (m) | Sampling Date | Volume Filtered (mL) | DNA Concentration (ng/µL) |
|---|---|---|---|---|---|
| TLW349 | Wishart | 3 | August 21, 2019 | 500 | 5.4 |
| TLW405 | Wishart | 0.76 (Under Ice) | January 22, 2020 | 1000 | 2.2 |

**Table S3** Sampling conditions for sample collection as prepared by Environment and Climate Change Canada and Natural Resources Canada field technicians. Conditions not provided for dates sampled are not available due to missing field data sheets.

| Lake | Sampling Date | Sampling Time | Cloud Coverage (%) | Wind | Air Temperature | Ice Cover |
|---|---|---|---|---|---|---|
| Wishart | July 16, 2018 | 2 :16 P.M. | 0 | Light | 24.3 | N/A |
| Little Turkey | July 16, 2018 | 10 :00 A.M. | 0 | Very Light | 17.5 | N/A |
| Big Turkey | July 18, 2018 | 11 :15 A.M. | 10 | Light | 23.4 | N/A |
| Wishart | August 14, 2018 | 11 :00 A.M. | 5 | Calm | 28.3 | N/A |
| Little Turkey | August 13, 2018 | 12 :40 P.M. | 0 | Light | 28.9 | N/A |
| Big Turkey | August 13, 2018 | 10 :15 A.M. | 0 | Light | 24.1 | N/A |
| Wishart | October 24, 2018 | 10 :15 A.M. | 10 | Light | 6.8 | Ice cover in AM |
| Little Turkey | October 25, 2018 | 12 :55 P.M. | 100 | Light | 4.8 | N/A |
| Big Turkey | October 25, 2018 | 10 :20 A.M. | 100 | Light | 5.4 | N/A |
| Little Turkey | June 28, 2019 | 12 :11 P.M. | 25 | Moderate | 26.4 | N/A |
| Big Turkey | June 28, 2019 | 9 :39 A.M. | 95 | None | 19.3 | N/A |
| Wishart | July 25, 2019 | 9 :40 A.M. | 0 | None | 21.2 | N/A |
| Little Turkey | July 24, 2019 | 12 :45 P.M. | 5 | Light | 23.1 | N/A |
| Big Turkey | July 24, 2019 | 10 :17 A.M. | 95% | None | 20.0 | N/A |
| Wishart | August 22, 2019 | 12 :30 P.M. | 65 | Light | 19.9 | N/A |
| Little Turkey | August 21, 2019 | 1 :00 P.M. | 40 | Moderate | 21.5 | N/A |
| Big Turkey | August 21, 2019 | 10 :37 A.M. | 10 | Moderate | 21.2 | N/A |

**Table S4** Probability values for cyanobacterial enrichment within the randomized total phylum counts per sample across a seasonal series. The mean and standard deviation of the randomized counts were used to calculate Z-scores. From the Z-scores, probability values were obtained and adjusted with a Bonferroni correction.

| Lake | Sampling Date | Mean | Standard Deviation | Observed | p-value | Adjusted p-value |
|---|---|---|---|---|---|---|
| Big Turkey | 20-Jan | 0.04 | 0.03 | 0.00 | 0.08 | 1.00 |
| Big Turkey | 19-Feb | 0.04 | 0.02 | 0.00 | 0.03 | 0.96 |
| Big Turkey | 19-Mar | 0.04 | 0.02 | 0.00 | 0.03 | 1.00 |
| Big Turkey | 19-May | 0.04 | 0.02 | 0.11 | 0.00 | 0.03 |
| Big Turkey | 19-Jun | 0.04 | 0.05 | 0.56 | 0.00 | 0.00 |
| Big Turkey | 18-Jul | 0.04 | 0.03 | 0.27 | 0.00 | 0.00 |
| Big Turkey | 19-Jul | 0.04 | 0.04 | 0.37 | 0.00 | 0.00 |
| Big Turkey | 18-Aug | 0.04 | 0.03 | 0.12 | 0.00 | 0.10 |
| Big Turkey | 19-Aug | 0.04 | 0.03 | 0.30 | 0.00 | 0.00 |
| Big Turkey | 18-Oct | 0.04 | 0.02 | 0.10 | 0.00 | 0.03 |
| Little Turkey | 20-Jan | 0.04 | 0.02 | 0.00 | 0.05 | 1.00 |
| Little Turkey | 19-Feb | 0.04 | 0.03 | 0.00 | 0.07 | 1.00 |
| Little Turkey | 19-Mar | 0.04 | 0.03 | 0.00 | 0.10 | 1.00 |
| Little Turkey | 19-May | 0.04 | 0.04 | 0.01 | 0.19 | 1.00 |
| Little Turkey | 19-Jun | 0.04 | 0.04 | 0.22 | 0.00 | 0.00 |
| Little Turkey | 18-Jul | 0.04 | 0.04 | 0.24 | 0.00 | 0.00 |
| Little Turkey | 19-Jul | 0.04 | 0.04 | 0.34 | 0.00 | 0.00 |
| Little Turkey | 18-Aug | 0.04 | 0.03 | 0.09 | 0.08 | 1.00 |
| Little Turkey | 19-Aug | 0.04 | 0.03 | 0.24 | 0.00 | 0.00 |
| Little Turkey | 18-Oct | 0.04 | 0.03 | 0.01 | 0.18 | 1.00 |
| Wishart | 20-Jan | 0.04 | 0.03 | 0.00 | 0.07 | 1.00 |
| Wishart | 19-Feb | 0.04 | 0.04 | 0.00 | 0.14 | 1.00 |
| Wishart | 19-Mar | 0.04 | 0.06 | 0.00 | 0.24 | 1.00 |
| Wishart | 19-May | 0.04 | 0.05 | 0.02 | 0.30 | 1.00 |
| Wishart | 19-Jun | 0.04 | 0.03 | 0.21 | 0.00 | 0.00 |
| Wishart | 18-Jul | 0.04 | 0.02 | 0.18 | 0.00 | 0.00 |
| Wishart | 19-Jul | 0.04 | 0.03 | 0.20 | 0.00 | 0.00 |
| Wishart | 18-Aug | 0.04 | 0.05 | 0.04 | 0.49 | 1.00 |
| Wishart | 19-Aug | 0.04 | 0.03 | 0.14 | 0.00 | 0.01 |
| Wishart | 18-Oct | 0.04 | 0.03 | 0.01 | 0.17 | 1.00 |

**Table S5** Probability values for cyanobacterial enrichment within the randomized total phylum counts per sample across a depth profile. The mean and standard deviation of the randomized counts were used to calculate Z-scores. From the Z-scores, probability values were obtained and adjusted with a Bonferroni correction.

| Lake | Month | Depth | Mean | Standard Deviation | Observed | p-value | Adjusted p-value |
|---|---|---|---|---|---|---|---|
| Big Turkey | August | Surface | 0.05 | 0.03 | 0.12 | 0.02 | 0.31 |
| Big Turkey | August | Secchi | 0.05 | 0.03 | 0.12 | 0.01 | 0.22 |
| Big Turkey | August | Deep | 0.05 | 0.04 | 0.13 | 0.03 | 0.45 |
| Big Turkey | July | Surface | 0.05 | 0.05 | 0.27 | 0.00 | 0.00 |
| Big Turkey | July | Secchi | 0.05 | 0.03 | 0.27 | 0.00 | 0.00 |
| Big Turkey | July | Deep | 0.05 | 0.06 | 0.45 | 0.00 | 0.00 |
| Little Turkey | August | Surface | 0.05 | 0.03 | 0.07 | 0.30 | 1.00 |
| Little Turkey | August | Secchi | 0.05 | 0.03 | 0.09 | 0.15 | 1.00 |
| Little Turkey | August | Deep | 0.05 | 0.04 | 0.13 | 0.02 | 0.45 |
| Little Turkey | July | Surface | 0.05 | 0.04 | 0.14 | 0.02 | 0.30 |
| Little Turkey | July | Secchi | 0.05 | 0.04 | 0.24 | 0.00 | 0.00 |
| Little Turkey | July | Deep | 0.05 | 0.05 | 0.14 | 0.03 | 0.53 |
| Wishart | August | Surface | 0.05 | 0.05 | 0.05 | 0.46 | 1.00 |
| Wishart | August | Secchi | 0.05 | 0.06 | 0.04 | 0.42 | 1.00 |
| Wishart | August | Deep | 0.05 | 0.03 | 0.05 | 0.47 | 1.00 |
| Wishart | July | Surface | 0.05 | 0.03 | 0.17 | 0.00 | 0.00 |
| Wishart | July | Secchi | 0.05 | 0.02 | 0.18 | 0.00 | 0.00 |
| Wishart | July | Deep | 0.05 | 0.02 | 0.16 | 0.00 | 0.00 |

**Table S6** Taxonomic classifications of cyanobacterial amplicon sequence variants performed with a Naïve-Bayes classifier trained on SILVA138 and confirmed with phylogenetic placement using Cydrasil.

| ASV Identifier | Classified Taxonomic Order |
|---|---|
| ASV475 | Sericytochomatia |
| ASV476 | Sericytochromatia |
| ASV650 | Oscillatoriales |
| ASV838 | Nostocales |
| ASV839 | Nostocales |
| ASV840 | Nostocales |
| ASV841 | Nostocales |
| ASV842 | Nostocales |
| ASV843 | Nostocales |
| ASV844 | Nostocales |
| ASV845 | Nostocales |
| ASV846 | Nostocales |
| ASV847 | Nostocales |
| ASV848 | Nostocales |
| ASV849 | Nostocales |
| ASV850 | Synechococcales |
| ASV851 | Synechococcales |
| ASV852 | Synechococcales |
| ASV853 | Synechococcales |
| ASV854 | Synechococcales |
| ASV855 | Synechococcales |
| ASV856 | Synechococcales |
| ASV857 | Synechococcales |
| ASV858 | Synechococcales |
| ASV859 | Synechococcales |
| ASV860 | Synechococcales |
| ASV861 | Synechococcales |
| ASV862 | Synechococcales |
| ASV863 | Synechococcales |
| ASV864 | Synechococcales |
| ASV865 | Synechococcales |
| ASV866 | Synechococcales |
| ASV867 | Synechococcales |
| ASV868 | Synechococcales |
| ASV869 | Synechococcales |
| ASV870 | Synechococcales |
| ASV871 | Synechococcales |
| ASV872 | Synechococcales |

| | |
|---|---|
| ASV873 | Synechococcales |
| ASV874 | Synechococcales |
| ASV875 | Synechococcales |
| ASV876 | Synechococcales |
| ASV877 | Synechococcales |
| ASV878 | Synechococcales |
| ASV879 | Synechococcales |
| ASV880 | Synechococcales |
| ASV881 | Synechococcales |
| ASV882 | Synechococcales |
| ASV883 | Synechococcales |
| ASV884 | Synechococcales |
| ASV885 | Synechococcales |
| ASV886 | Synechococcales |
| ASV887 | Synechococcales |
| ASV889 | Synechococcales |
| ASV890 | Synechococcales |
| ASV891 | Synechococcales |
| ASV892 | Synechococcales |
| ASV893 | Synechococcales |
| ASV894 | Synechococcales |
| ASV895 | Synechococcales |
| ASV1166 | Synechococcales |
| ASV1167 | Synechococcales |
| ASV1168 | Synechococcales |
| ASV1169 | Synechococcales |
| ASV1170 | Synechococcales |
| ASV1171 | Synechococcales |
| ASV1172 | Chroococcales |
| ASV1174 | Chroococcales |
| ASV1175 | Chroococcales |
| ASV1176 | Chroococcales |
| ASV1177 | Chroococcales |
| ASV1178 | Chroococcales |
| ASV1179 | Chroococcales |
| ASV1180 | Chroococcales |
| ASV1181 | Chroococcales |
| ASV1182 | Chroococcales |
| ASV1183 | Chroococcales |
| ASV1184 | Chroococcales |
| ASV1185 | Synechococcales |
| ASV1186 | Synechococcales |
| ASV1187 | Synechococcales |
| ASV1188 | Synechococcales |
| ASV1189 | Synechococcales |

| | |
|---|---|
| ASV1190 | Chroococcales |
| ASV1191 | Chroococcales |
| ASV1192 | Chroococcales |
| ASV1193 | Synechococcales |
| ASV1194 | Chroococcales |
| ASV1195 | Chroococcales |
| ASV1197 | Spirulinales |
| ASV1199 | Chroococcales |
| ASV1200 | Chroococcales |
| ASV1201 | Chroococcales |
| ASV1202 | Chroococcales |
| ASV1203 | Chroococcales |
| ASV1204 | Chroococcales |
| ASV1205 | Chroococcales |
| ASV1207 | Chroococcales |
| ASV1208 | Chroococcales |
| ASV1209 | Chroococcales |
| ASV1210 | Chroococcales |
| ASV1211 | Chroococcales |
| ASV1212 | Synechococcales |
| ASV1213 | Synechococcales |
| ASV1214 | Synechococcales |
| ASV1215 | Synechococcales |
| ASV1216 | Synechococcales |
| ASV1217 | Synechococcales |
| ASV1218 | Synechococcales |
| ASV1219 | Synechococcales |
| ASV1220 | Synechococcales |
| ASV1221 | Oscillatoriales |
| ASV1223 | Chroococcales |
| ASV1224 | Synechococcales |
| ASV1225 | Synechococcales |
| ASV1226 | Oscillatoriales |
| ASV1227 | Oscillatoriales |
| ASV1228 | Chroococcales |
| ASV1229 | Chroococcales |
| ASV1230 | Gloeobacterales |
| ASV1231 | Gloeobacterales |
| ASV1232 | Candidatus Melainabacteria |
| ASV1233 | Candidatus Melainabacteria |
| ASV1234 | Candidatus Melainabacteria |
| ASV1235 | Candidatus Melainabacteria |
| ASV1236 | Candidatus Melainabacteria |
| ASV1237 | Candidatus Melainabacteria |
| ASV1238 | Candidatus Melainabacteria |

| | |
|---|---|
| ASV1239 | Candidatus Melainabacteria |
| ASV1240 | Candidatus Melainabacteria |
| ASV1241 | Candidatus Melainabacteria |
| ASV1242 | Candidatus Melainabacteria |
| ASV1243 | Candidatus Melainabacteria |
| ASV1244 | Candidatus Melainabacteria |
| ASV1245 | Candidatus Melainabacteria |
| ASV1246 | Candidatus Melainabacteria |
| ASV1247 | Candidatus Melainabacteria |
| ASV1248 | Candidatus Melainabacteria |
| ASV1249 | Candidatus Melainabacteria |

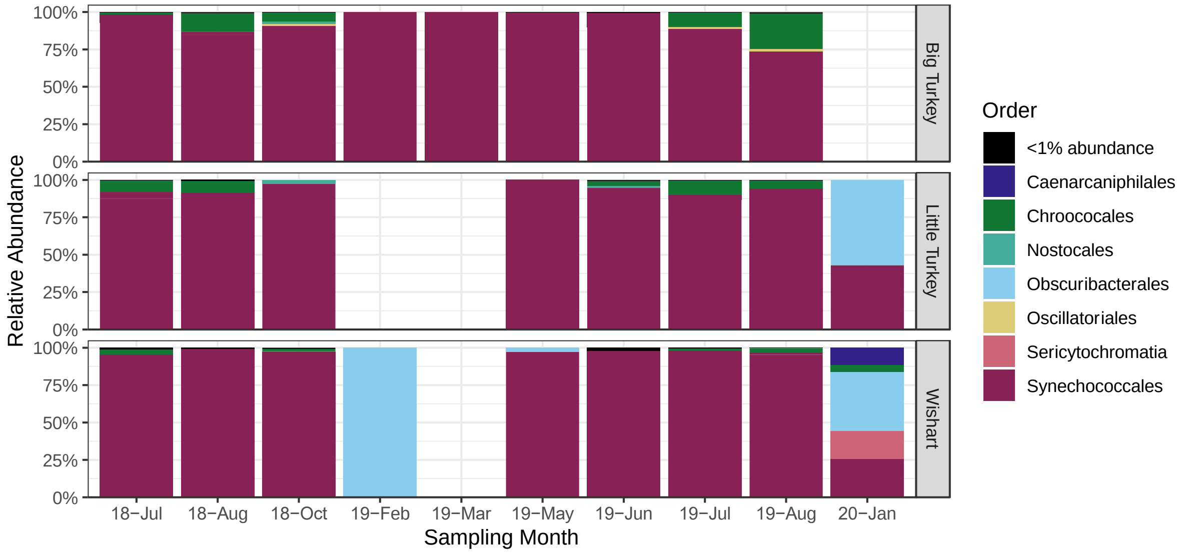

**Figure S4** Stacked bar charts depicting the relative abundances of cyanobacterial families composing the cyanobacterial communities identified from amplicon sequencing of the V4 region of the 16S rRNA gene across a seasonal sampling series in a shallow lake (Wishart), mid-sized lake (Little Turkey) and deep lake (Big Turkey). Notably, cyanobacteria were present in less than 1% abundance in the bacterial communities during ice-covered months. Sampling months with no bar signify the absence of cyanobacterial sequences.

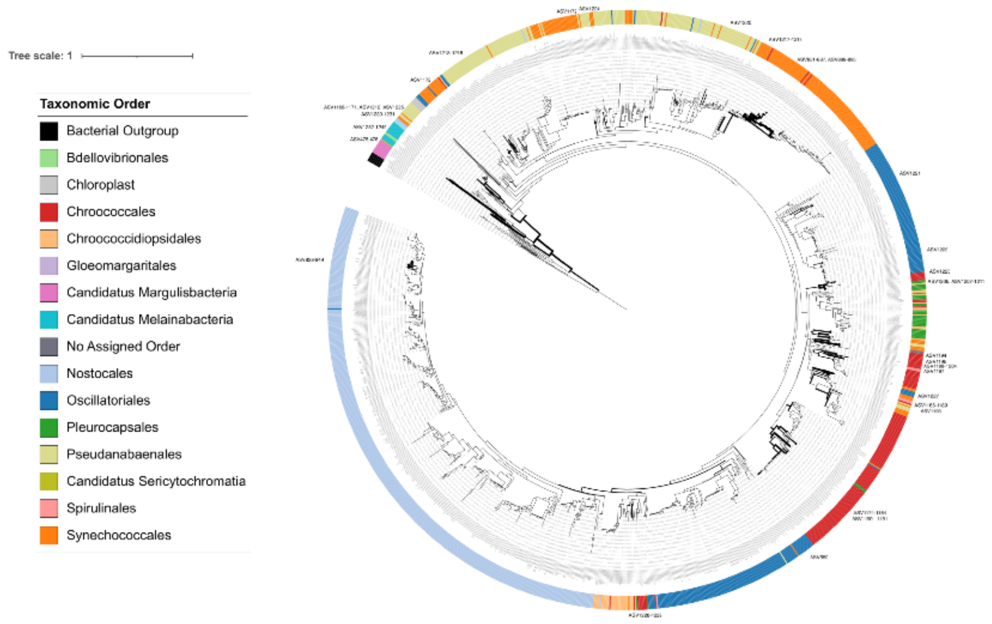

**Figure S5** Phylogenetic tree indicating the taxonomic affiliation of amplicon sequence variants displayed using the Interactive Tree of Life. Cydrasil was used to place cyanobacterial amplicon sequence variants onto a curated cyanobacterial 16S rRNA reference tree. Taxonomic orders of references are indicated by the coloured strip. Thickness of branches on the tree indicate the placement locations of query amplicon sequence variants.

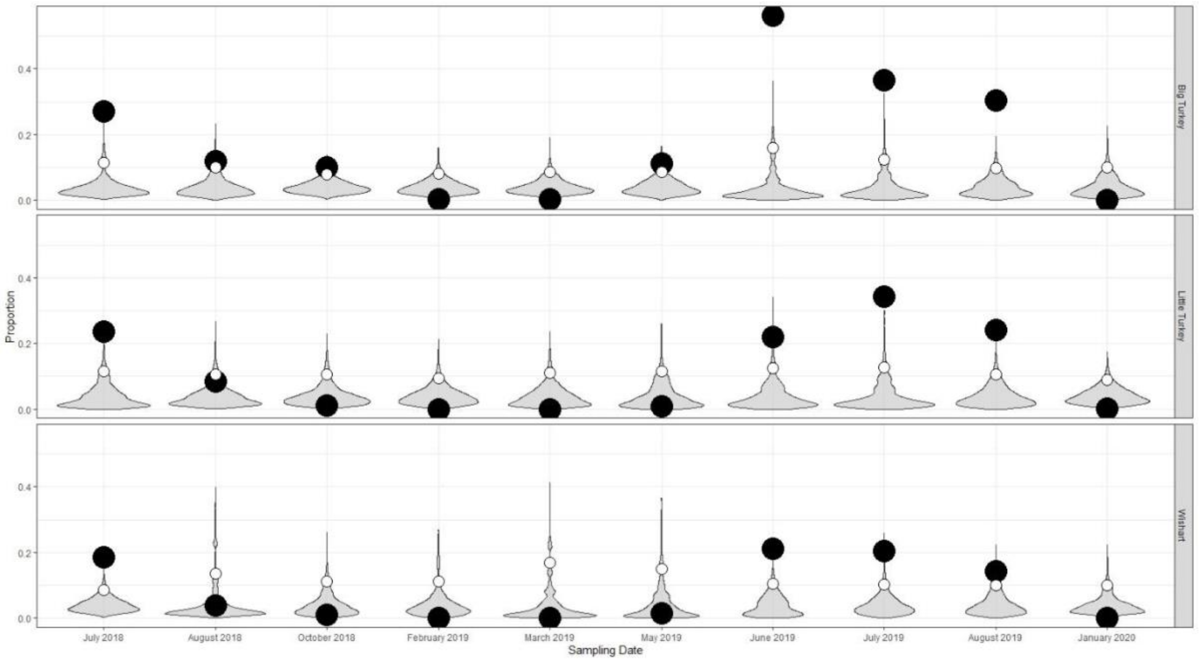

**Figure S6** Violin plots visualizing the distribution of randomized total phylum counts and the specific relative abundance of cyanobacteria relative to the 95$^{th}$ percentile across sampling months in a multi-seasonal period in Big Turkey, Little Turkey and Wishart Lake. The observed cyanobacterial values are visualized with the solid black circle, and the 95$^{th}$ percentile with the white circle. Black dots present above the white circle represents an enriched abundance of cyanobacteria present in the bacterial community.

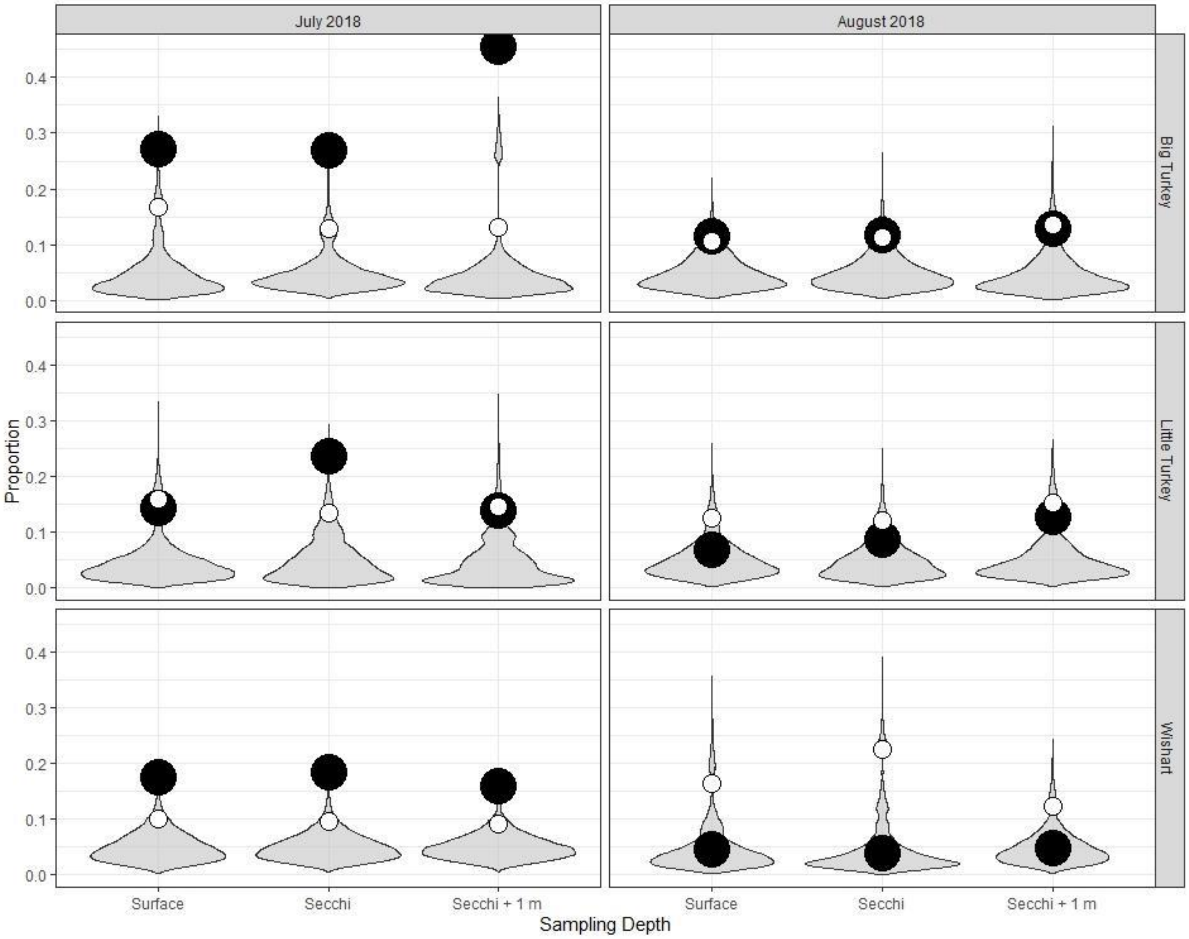

**Figure S7** Violin plots visualizing the distribution of randomized total phylum counts and the specific relative abundance of cyanobacteria relative to the 95$^{th}$ percentile across sampling depths in Big Turkey, Little Turkey and Wishart Lake. The observed cyanobacterial values are visualized with the solid black circle, and the 95$^{th}$ percentile with the white circle. Black dots present above the white circle represents an enriched abundance of cyanobacteria present in the bacterial community.
